# Evolutionary dynamics of genome structure and content among closely related bacteria

**DOI:** 10.1101/2024.07.08.602537

**Authors:** Marco Molari, Liam P. Shaw, Richard A. Neher

## Abstract

Bacterial genomes primarily diversify via gain, loss, and rearrangement of genetic material in their flexible accessory genome. Yet the dynamics of accessory genome evolution are very poorly understood, in contrast to the core genome where diversification is readily described by mutations and homologous recombination. Here, we tackle this problem for the case of very closely related genomes. We comprehensively describe genome evolution within n=222 genomes of *E. coli* ST131, which likely shared a common ancestor around one hundred years ago. After removing putative recombinant diversity, the total length of the phylogeny is 6000 core genome mutations. Within this diversity, we find 22 modifications to core genome synteny and estimate around 2000 structural changes within the accessory genome, i.e. one structural change for every 3 core genome mutations. 63% of loci with structural diversity could be resolved into individual gain and loss events with ten-fold more gains than losses, demonstrating a dominance of gains due to insertion sequences and prophage integration. Our results suggest the majority of synteny changes and insertions in bacterial genomes are likely deleterious and only persist for a short time before being removed by purifying selection.

## I. INTRODUCTION

Many microbial species are able to acquire new genes and advantageous alleles in a process known as Horizontal Gene Transfer (HGT) (Arnold *et al*., 2022), in which genetic material is transferred from one microbe to the next. The horizontal movement of genes is often mediated by Mobile Genetic Elements (MGEs) (Haudiquet *et al*., 2022; Tokuda and Shintani, 2024) such as plasmids, transposons and phages. These elements participate in a complex evolutionary interplay with their host and sometimes carry cargo that is beneficial to the host, such as antibiotic resistance genes or defense systems against other MGEs (Georjon and Bernheim, 2023; Mayo-Muñoz *et al*., 2023). This makes HGT one of the major drivers of the spread of antimicrobial resistance (Partridge *et al*., 2018; Von Wintersdorff *et al*., 2016), and understanding its dynamics and effects on the microbial genome is of crucial importance to combat this spread.

As a consequence of HGT, bacteria in the same species can differ dramatically in the content of their genomes, with any given individual only possessing a fraction of the total collection of genes present in the species, named the *pangenome* (Brockhurst *et al*., 2019). The pangenome can be split in two components: the *core* genome, which refers to the set of genes that are common to all isolates, and the *accessory* genome, indicating genes that are specific to a subset of isolates, sometimes associated to adaptations to specific niches or lifestyles (Touchon *et al*., 2020).

Given this split into core and accessory genome, the similarity of genomes is often quantified by two different metrics: the number of mutations in the alignable core regions and the overlap within the accessory genome – that is, how many accessory genes two genomes have in common. In many microbial species, genomes that differ by very few mutations in their core genome can show large differences in accessory genome content (Doolittle and Zhaxybayeva, 2009; Touchon *et al*., 2009, 2020). The horizontal exchange of accessory genes thus has to be very fast, but a quantitative understanding of this dynamics is lacking. Yet despite this flexibility the structural organization of the genome remains strongly conserved, with the order of core genes being maintained even across large evolutionary distances (Rocha, 2006, 2008).

To quantify how genomic diversity is generated and accumulates over time, one would need to observe individual structural modifications on short evolutionary timescales. However, systematically identifying structural changes (gain, loss or changes in gene order) and quantifying their rates is a difficult problem with two complementary challenges. First, detecting structural changes across a large set of isolates requires comparisons of only-partially-alignable and potentially shuffled portions of the genome, which is technically and computationally challenging. Second, homologous recombination in the core genome is sufficiently rapid that the genealogy of bacteria can often no longer be reconstructed (Sakoparnig *et al*., 2021), meaning one lacks the consistent estimates of divergence between strains which are necessary to calibrate rates of structural evolution.

Here, we aim to overcome these challenges and accurately quantify rates of structural evolution in *E. coli*. To address the challenge of a reliable clonal phylogeny, we focus on the recently-expanded sequence type ST131 where homologous recombination is still a perturbation that can be controlled for. To identify individual modifications to genome structure and content, we use a *pangenome graph* representation for the genomes in our dataset, as provided by *PanGraph* (Noll *et al*., 2023). Within this gene-agnostic representation, genomes are encoded as paths through alignments of homologous regions. Any structural changes appear as deviations between paths. This approach allows us to identify changes in core genome order and find regions harbouring structural changes in the accessory genome. We can then not only quantify their diversity and size but also infer the rates at which structural diversity accumulates, thereby providing a quantitative picture of the evolution of genome structure on short timescales.

## II. DATASET OVERVIEW AND GRAPH CONSTRUCTION

The multidrug-resistant sequence type ST131, first identified in 2008, is associated with the expression of extended-spectrum beta-lactamases (ESBLs) and is a major cause of antibiotic-resistant urinary tract infections worldwide (Decano and Downing, 2019; Pitout and Finn, 2020; Stoesser *et al*., 2016). The most recent common ancestor of all ST131 is estimated to have lived around 1900 (95% HPD 1842–1948 (Ludden *et al*., 2020)) with most of the clonal expansion taking place in the last 35 years driven by the widespread use of beta-lactam antibiotics (Ben Zakour *et al*., 2016; Gladstone *et al*., 2021; Kallonen *et al*., 2017; Stoesser *et al*., 2016). Due to its importance for public health there are a large number of high-quality *E. coli* ST131 genomes in public databases, representing a set of closely related genomes that we can be confident diverged only recently.

We downloaded all complete *E. coli* assemblies from RefSeq (O’Leary *et al*., 2016), and selected the ones that were classified as ST131 by multi-locus sequence typing and had a mash (Ondov *et al*., 2016) distance *<* 0.008 from the ST131 reference `NC 013654.1` (see *SI* sect. 1 for more details). After quality control and filtering the dataset included *n* = 222 isolates (average genome size 5.1*±*0.1 Mbp), with a large number of genomes from environmental sampling in Switzerland (Biggel *et al*., 2023) and hospital sampling in Sweden (Jaén-Luchoro *et al*., 2023) (see *SI* fig. 1). We used PanGraph (Noll *et al*., 2023) to losslessly encode all chromosomal sequences in the form of a pangenome graph (a ‘pangraph’) that we could query for all downstream analyses (see *SI* sect. 2). By summing the lengths of all block sequences in our graph we get a total pangenome size of 8 Mbp. Analogously, the core-genome size is 3.6 Mbp and the soft-core (present in *>* 95% of isolates) 4.2 Mbp (see *SI* fig. 3).

**FIG. 1.**
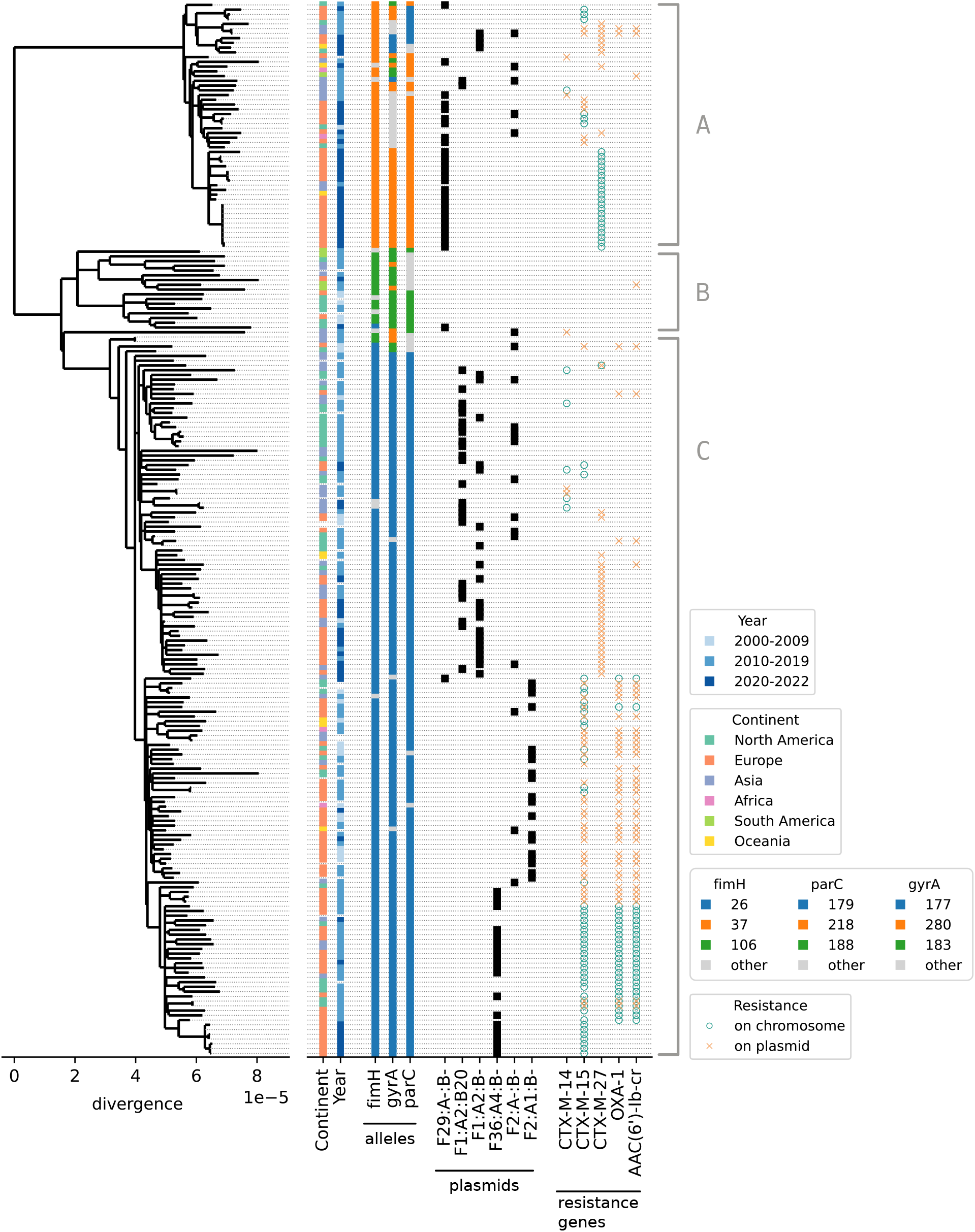
Core genome tree reconstructed after removing regions with suspected recombination. Annotations on the right report geographic origin and isolation year, *fimH, gyrA* and *parC* alleles, IncF plasmid type, and presence of particular resistance genes, see *SI* sect. 1 for details. The tree recapitulates the known global diversity of *E. coli* ST131 with representatives from the tree main clades A, B and C.

**FIG. 2.**
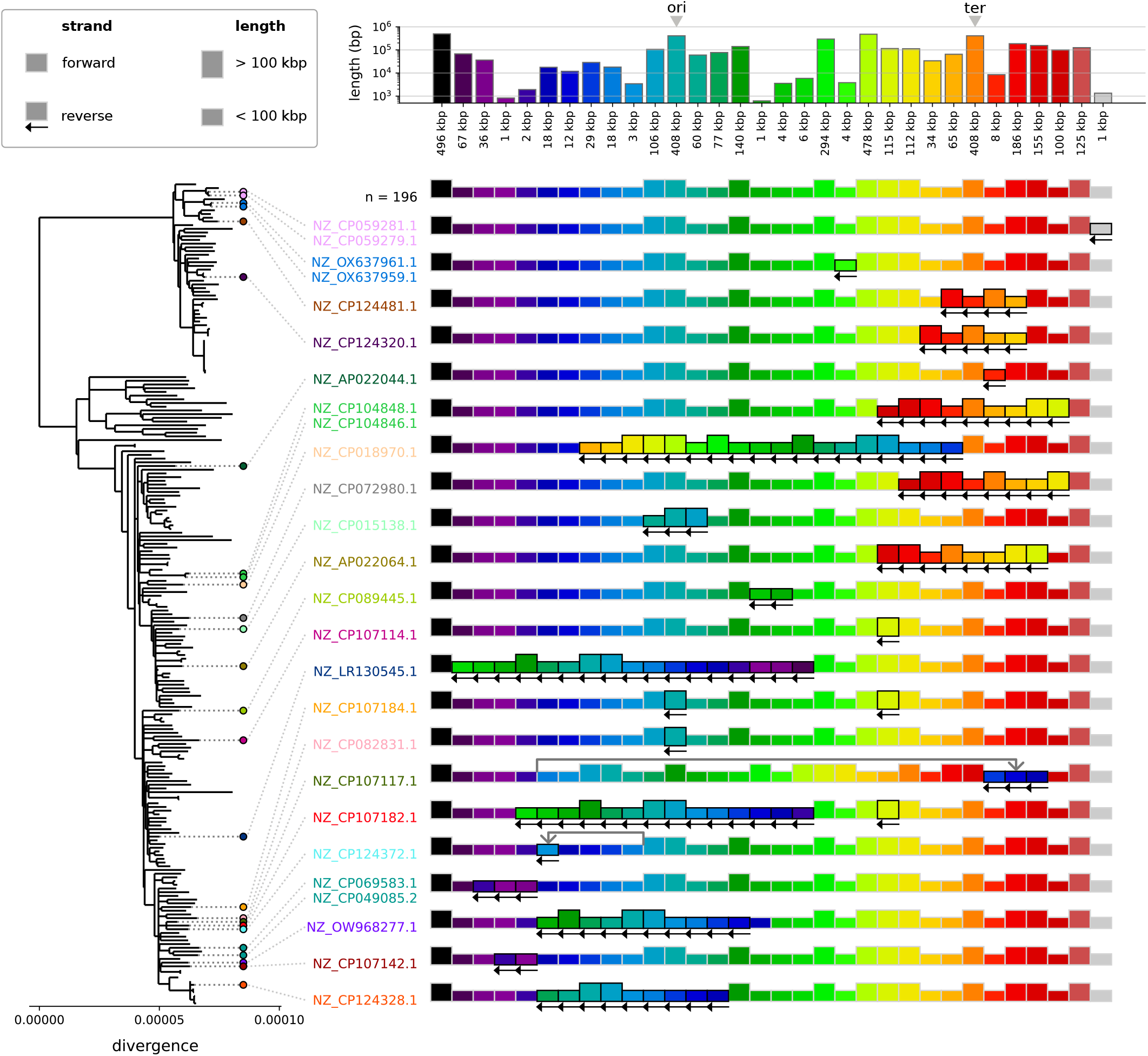
Variation in core genome synteny across ST131. Each block represents a region of core sequence that is syntenic in all isolates, and each row a particular order of these regions in the chromosome. Most isolates (196/222) have the consensus order reported in the first row. Each subsequent row represents a different order present in other isolates, highlighted in corresponding color on the core genome tree. The height of a block is related to its length (see top), while left-pointing arrows indicate reverse-complemented blocks. Note that this analysis uses the entire core genome (3.59Mbp) before filtering out suspected homologous recombination.

**FIG. 3.**
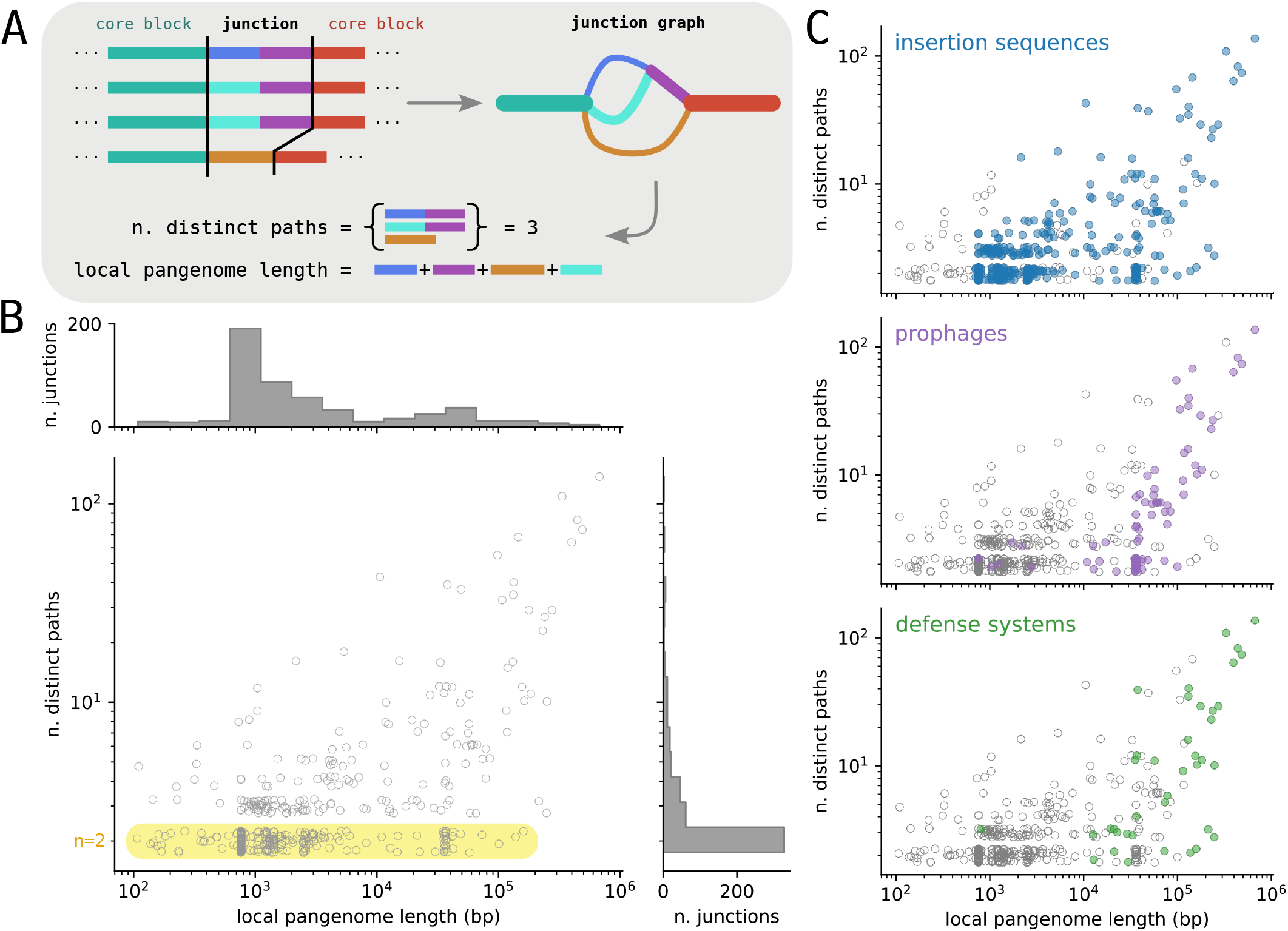
Structural diversity in the accessory genome. **A**: Junction graphs are created by running PanGraph on a region spanning the interval between two consecutive core blocks, including the core blocks themselves. For each graph we evaluate the number of different distinct paths and the total local accessory pangenome length. **B**: Joint distribution of junction size, measured as accessory pangenome content, and complexity, measured as total different number of paths, for the backbone junctions in our dataset. To better visualize regions of high density where many points are overlapping, we added a vertical uniform random displacement to each point of magnitude *±*0.25. Binary junctions with only two alternative paths are highlighted in yellow. Panel **C** shows the same distribution as panel B and highlights junctions that contain insertion sequences (blue), prophages (purple) and defense systems (green). See main text for details.

As expected, isolates from the same sequence type are extremely similar in their core genome (less than 0.01% divergence on average if we exclude recombined regions, see below). However, they can differ by hundreds of kbp in their accessory genome (see *SI* fig. 6).

As a first step, we extracted all core blocks from the pangenome graph and built a core genome alignment. Due to the recent divergence of ST131 the density of SNPs in this alignment is very small, except for regions that underwent homologous recombination with isolates outside of ST131. These regions are visible as islands of high SNP density (Sakoparnig *et al*., 2021) (see *SI* fig. 4). Using a strategy similar to Gubbins (Croucher *et al*., 2015), we excluded these regions by filtering out from the alignment any 2kbp window in which an isolate has more than 3 mutations with respect to the consensus (see *SI* sect. 3). This filtering procedure reduced the size of the core genome alignment roughly by 1/3 (from 3.6 Mbp to 2.4 Mbp) while at the same time removing 70% of the polymorphic positions (from 22k to 6k polymorphic sites in the alignment) that are clustered in the recombined regions.

**FIG. 4.**
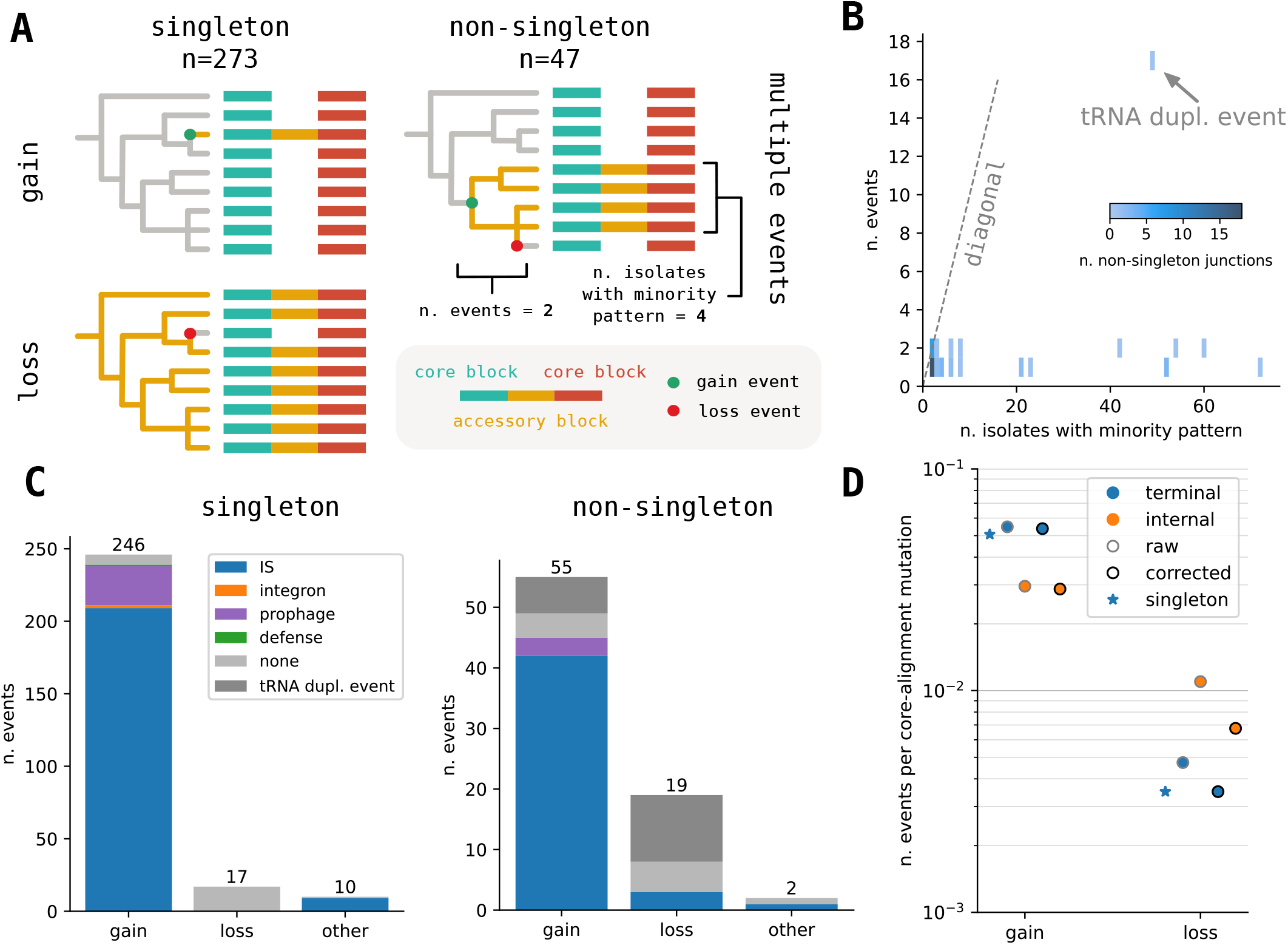
Quantifying rates of genome structure evolution. **A**: Junctions with *singleton* structural diversity are trivially classified as single gain or less events. *Non-singleton* junctions, i.e. where more than one isolate follows the majority path, are often also explained by a single event, but can have more complex pattern and require multiple gain/loss events to be explained. **B**: The number of gain/loss events required (y-axis) to explain non-singleton junctions is mostly one or two regardless of the prevalence of the minority path (x-axis), with a single exception that corresponds to a tRNA duplication event. **C**: number of gain/loss/other events for singleton and non-singleton junctions, stratified by associated annotation (see main text). **D**: Estimates of gain and loss rates obtained by dividing the number of events by the corresponding length of the core genome tree measured as expected number of mutations. The different estimate are for: just singleton events (star), all events on terminal branches (blue) and events on internal branches (orange). The correction (black edge) corresponds to removing events attributed to the single junction featuring the tRNA duplication.

After removing these putative recombinant regions, we used *FastTree* (Price *et al*., 2010) to infer a phylogenetic tree from the filtered alignment (fig. 1). The resulting phylogeny is compatible with the known global diversity of *E. coli* ST131, with representatives from the three main clades A, B and C (Pitout and Finn, 2020). Locus typing, plasmid typing, and resistance gene presence are broadly consistent with the phylogeny. This phylogeny and the filtered alignment are mostly compatible and only a few homoplasies remained when inferring ancestral sequences on the phylogeny (2.8% of all polymorphic sites, compared to 35% before recombination filtering, see *SI* sect. 3B and *SI* fig. 5). This suggests that our recombination filtering not only removed recombination with divergent genomes outside of our dataset, but that residual recombination within our dataset does not substantially distort the phylogeny.

## III. CORE GENOME SYNTENY MUTATIONS

Previous work has established that the order of core genes is conserved even across large evolutionary distances (Rocha, 2006, 2008; Touchon *et al*., 2009). Yet it is unclear whether this conservation is due to a very low rate of introduction of synteny changes (“synteny mutation rate”), or because such changes to synteny are deleterious and never persist. We therefore first surveyed patterns of synteny variation in our *E. coli* ST131 dataset. Considering only alignable regions (blocks) larger than 500bp, the 3.59Mbp core genome of ST131 is divided into 32 maximal syntenic regions (range: 1 − 478kbp; see *SI* sect. 3.C).

The vast majority of isolates in our dataset (196/222) have these blocks in the same consensus order and the remaining 26 isolates show 22 different synteny patterns (fig. 2; see *SI* fig. 7 for the consensus order mapped to the genome). Most of the non-consensus patterns are observed in a single isolate, likely caused by events that occurred on the terminal branches of the phylogenetic tree. There are however 4 patterns that are shared by pairs of isolates in close phylogenetic proximity, indicating that the patterns likely originated from a single event on an internal branch of the tree. Most patterns deviate from the consensus pattern by a single inversion, often centered around the origin or terminus of replication consistent with previous observations (Eisen *et al*., 2000). There are four exceptions: the patterns of isolates `NZ CP107184` and `NZ CP107182` feature two inversions, while in isolates `NZ CP107117` and `NZ CP124372` the inverted region is also translocated.

Isolates `NZ CP069583.1` and `NZ CP049085.2` are the only case of a synteny pattern shared by two isolates that are not nearest neighbors. These form a clade together with isolate `NZ CP103557.1`, which does not feature the inversion. We investigated this case in greater detail (see *SI* fig. 8) and found a high density of repetitive sequence flanking the inversion, compatible with a reversion of this inversion.

The total tree length corresponds to 6040 mutations in the core genome, implying that events that change core genome synteny happen at a rate of roughly 1 one every 270 mutations on the (filtered) core genome. Note that here we only consider clonally acquired mutations and exclude the ones likely obtained in recombination events. The fact that the majority of synteny changes is observed on terminal branches suggest that many of them might be deleterious. However, as expected for populations that recently expanded, the core genome tree is dominated by terminal branches (80% of total branch length) such that there is little power to test for over-representation of inversions or translocations on terminal branches.

## IV. ACCESSORY GENOME DIVERSIFICATION

The strong conservation of synteny in ST131 provides a good frame of reference to study variation of the accessory genome. Any pair of core blocks defines a specific context that is present in all isolates with very few exceptions due to synteny breaks (see *SI* fig. 10). Accessory genome variation between a pair of consecutive core blocks can thus be meaningfully compared across isolates in the dataset. This diversity likely arose through gains and loss since the most recent common ancestor of the genomes in the dataset.

We call a pair of consecutive core blocks and the enclosed accessory genome a core genome *junction*. We analyzed each junction in isolation by extracting the genetic region that spans the two flanking core blocks from every isolate and then use PanGraph to build a small local genome graph for this region (fig. 3A), related to the idea of local graphs in (Colquhoun *et al*., 2021). While this local graph is in principle embedded in the global graph, rebuilding the local graph increases reliability and avoids excessive graph fragmentation due to homology with other regions in the genome, e.g. for heavily duplicated elements. We found a total of *n* = 519 such core block pairs flanking accessory variation, for details see *SI* sect. 6.

We characterized each junction by its *diversity*, measured in terms of number of distinct realized paths between the flanking core blocks, and its *size*, expressed as the total length (bp) of the accessory genome contained between the two core blocks (*local pangenome length)*. The joint distribution of these two quantities (fig. 3B) shows the large heterogeneity of structural diversity across the genome. 328 of the 519 junctions (63%) display only two different distinct paths, and often one of the two is an empty path (i.e. no accessory content) correspond to an individual gain/loss event. Their size distribution is bimodal, with a peak around 1 kbp due mainly to insertion sequences, and another peak at 30-40 kbp corresponding to prophages (fig. 3C). On the other end of the spectrum are a small number of junctions of very high diversity, with many different paths and more than 100kbp of distinct genetic sequence.

We characterized junctions by the presence of specific features such as insertion sequences (IS), prophages, or defense systems (see fig. 3C). IS are detected using *ISEScan* (Xie and Tang, 2017). Each isolate has an average of 70 ISs on their chromosome. The three most represented families are IS3, IS66 and IS21, see *SI* fig. 11. ISs are present in most junctions (451/519) and account for the large number of binary junctions of size 1 kbp. Prophages, detected using *geNomad* (Camargo *et al*., 2023), are present in many large and complex junctions, and they also account for a large fraction of binary junctions of size 30-40 kbp (90% of such junctions contain prophages). Each chromosome harbors on average 6 prophages (*SI* fig. 12). Defense systems were detected using *DefenseFinder* (Tesson *et al*., 2022). As expected they are located mostly in large and diverse junctions and are often carried by prophages or co-localized with insertion sequences. Each isolate has around 10 different defense systems (*SI* fig. 13). We also verified that most of the diverse junctions identified by our analysis are in the same core genome location as the defense system ‘hotspots’ identified in (Hochhauser *et al*., 2023) (see *SI* sect. 10 and *SI* fig. 17). Finally, we used *Integron-Finder* (Néron *et al*., 2022) to locate integrons. However we could detect only 11 integron-associated junctions in our graph.

## V. ESTIMATING RATES OF ACCESSORY GENOME EVOLUTION

The structure of many of the diverse junctions with their dozens of different paths is hard to decompose into individual gain, loss, or transposition events. We therefore first focus on the 63% of junctions with binary variation (fig. 3B) whose diversity can usually be interpreted in terms of gain or loss of accessory segments.

We only considered binary *backbone* junctions (n=320), i.e. junctions associated to core-edges that are present in all isolates. This only excluded few (n=8) junctions flanking synteny breaks in some isolates. We further subdivided these junctions into *singletons* (n=273, 85%), i.e. where the minority pattern is present only in one isolate, and *non-singletons* (n=47, 15%) (fig. 4A). Determining whether a singleton is a gain or loss event amounts to verifying whether the accessory region in the minority pattern is a strict subset (*loss*) or superset (*gain*) of the accessory region of the majority pattern. In case none of the two conditions was met (e.g. a mixture of loss and gain, an inversion or a translocation) we classified the event as *other*. We find that the majority of events are gains (n=246, 90%), and only a minority (n=17, 6%) losses (fig. 4C).

For non-singleton junctions we inferred a parsimonious set of gain and loss events to explain the presence/absence pattern of the two alleles across the phylogeny using TreeTime (Sagulenko *et al*., 2018) (fig. 4A). The inferred events can be on internal or terminal branches. Fig. 4B shows the distribution of number of events required to explain the pattern of each non-singleton junction, along with the number of isolates with the minority pattern. While a random pattern would require a number of events roughly equal to the number of isolates with the minority pattern, we found that all patterns can be explained with just one or two events, with the exception of a single junction. The latter is due to a tRNA duplication event that appears to have been lost multiple times in clade A; the presence/absence pattern requires 17 independent events. High rates of *pheV* tRNA insertion have been previously described by (Touchon *et al*., 2009), but a different tRNA is involved here (*Tyr* -tRNA). As before the great majority of events are gains, but losses are a 4-fold or 2-fold higher proportion for non-singleton than for singletons depending on whether or not the tRNA duplication event is counted, respectively.

To gain insight into the dominant mobilisation mechanisms, we tried to associate gain or loss events with particular MGEs using the annotations provided by the tools discussed above. When more than one signature MGE is present, we associated a junction with one category using assignment hierarchy: integrons *>* prophages *>* defense systems *>* ISs. This prioritizes larger or rarer elements. We found that the great majority of gains were associated with IS elements, followed by prophages (fig. 4C).

Having mapped all the events on the phylogeny where a junction became structurally polymorphic with two alleles, we can estimate the rate of such events in comparison to the number of core genome SNPs on the tree like we did above for modifications of core genome synteny. We estimate these rates separately for terminal and internal tree branches, which account for 80% and 20% of the total branchlength, respectively. Rates on terminal branches can be estimated either using junctions with *singleton* variation or including events from *non-singleton* junctions inferred to have happened on terminal branches. The two estimates are very similar, especially if the tRNA duplication event is discarded, and indicate that a gain event is detected every roughly 20 core genome mutations, while a loss event roughly every 300 (see fig. 4D). For internal branches, gain rates are slightly lower (one event per 35 core mutations) and loss rates are slightly higher (one event per 150 core mutations). We further confirmed that there is a noisy but clear correlation between the length a branch and the number of events mapped to it (*SI* fig. 14).

Our analysis reveals that ISs are responsible for most of the binary structural genomic diversity, and mostly in the form of insertions present in a minority of isolates. These insertions might have a fitness effect by disrupting functional genes. To investigate the patterns of gene disruption we considered all binary junctions with an IS gain, and checked the gene annotations spanning the break point in genomes that do not contain the IS (see *SI* sect. 9 *SI* fig. 15). In 71% of cases the IS interrupted a gene, thus resulting in a loss of a gene. If insertion happened uniformly across the genome, one would expect the fraction of gene-disrupting insertions to be similar to the fraction of 88.4% of the *E. coli* core genome that is coding sequence. The underrepresentation of gene-disrupting insertions is probably because of purifying selection. If we assume that insertions in non-coding regions have small fitness effects, while insertions in coding regions are on average deleterious, the observed excess of insertions in non-coding regions suggests that roughly 60% of gene-disrupting insertions are removed by purifying selection before being observed, and that the rate of gain events before selection is higher than our estimate. Note, however, that insertions in non-coding regions can have fitness effects as well, for example by carrying promoters that can activate expression of downstream genes (Siguier *et al*., 2014).

In contrast to binary junctions, for junctions with more than two alternative paths it is often impossible to decompose the diversity into individual events. Inspecting the most complex junctions suggests that they have almost certainly exchanged genetic material with a diverse reservoir many times. While we cannot estimate separate rates for gain and loss, we can nevertheless estimate a lower bound for the overall rate of structural diversification. If a junction has *n* realized paths, there must have been at least *n* − 1 structural modifications to generate this diversity from a single ancestral form. We thus summed *n*_*i*_ − 1 for all junctions *i* and thereby estimate that at least ∼2000 modifications have taken place in the evolutionary history of *E. coli* ST131 (Σ_*i*_ (*n*_*i*_−1) = 1936, see *SI* fig. 23) – about one for every three core genome mutations. One sixth of these structural modifications fall into the 320 binary junctions, and half of them into the 20 most complex junctions.

To illustrate the structural diversity found in complex junctions we picked two examples and created a linear representation for their structure and content. *SI* figs. 20 and 21 show the junction corresponding to *hotspot 18* in (Hochhauser *et al*., 2023). This hotspot is empty in all but one isolate of clade A, which harbors a prophage. A different prophage is integrated in clade B and C, with nested diversity due to the movement of ISs. Overall this junction features 33 different paths and a total accessory genome length of 107 kbp. *SI* figs. 18 and 19 show the junction corresponding to Hochhauser *hotspot 11*. Diversity in clade A is generated mainly by the integration of a large segment containing several ISs and defense systems. In clade B and C structural diversity is generated by the integration of different prophages and the movement of ISs. This junction features 27 different paths and a total pangenome length of 240 kbp. Overall visual inspection of the structural diversity within these regions reveals both large-scale differences, that are broadly compatible with the phylogenetic structure, but also nested small-scale changes generated by the movement of small segments. Note that in both examples, even when empty the hotspot region contains tRNA genes, which are known targets for integrase enzymes (Bellanger *et al*., 2014; Williams, 2002).

## VI. DISCUSSION

Analyses in molecular evolution are typically based on SNPs in alignments of homologous sequences. From such data, it is straightforward to define distances and, in case of asexual inheritance, reconstruct phylogenetic trees that approximate the evolutionary history of the population. We have a good understanding of the elementary evolutionary process – mutation – which generates this diversity. Mutation can be described by substitution models that define rates for transitions between its discrete states. The maturity of this field and the associated analysis tools is one of the reason why bacterial genomes are often analyzed in a similar way: identify the core genome and analyze SNPs in this core genome. However, two bacterial genomes that differ only at a few hundred positions in the core genome can differ by hundreds of thousands of bases in the accessory genome, and by focusing on the core genome most of the evolutionary dynamics are missed.

Developing a quantitative description of evolution of the accessory genome, analogous to that of mutations in alignments of homologous sequences, is a hard problem. Instead of mutating between a fixed set of nucleotides, the accessory genome evolves by gain of genetic material of various length from a diverse pool, loss of segments, inversions, translocations, duplications, etc. Furthermore, the rates of these processes vary dramatically and are unknown to many orders of magnitude. Here, we systematically surveyed all structural changes in a population of recently diverged bacteria using pangenome graphs to begin to quantify these rates.

We chose *E. coli* ST131 because its recent divergence means horizontal processes are rare enough that they can mostly be identified as single events on a robust core genome phylogeny. This then allowed us to estimate rates of the processes that change genome content and structure, at least to within an order of magnitude, by comparing the number of events to the length of the core genome tree. We found that core genome synteny changes at a rate of one event for every 250 mutations in the core genome, with the majority of changes being inversions. Extrapolation of this rate to typical core genome distances of two *E. coli* genomes results in around 100 synteny breaks, which is not observed (see *SI* fig. 9). This is consistent with the notion that microbial genomes are under selection to maintain their rigid organization (Rocha, 2008).

Our survey of variation in the accessory genome highlights the heterogeneity of genome structure evolution. We identified 519 loci with accessory genome variation with at least 2000 structural modifications to the genome, suggesting that one structural change occurs every three core genome mutations. At around 300 of these loci, we found only two distinct structural variants which allows to decompose the diversity into individual gain and loss events. From these region, we estimated that gain of genetic material (insertion) is about 10 times more frequent than loss. Despite overall low diversity of ST131, we found a multitude of different variants often consisting of multiple MGEs at about 200 other genomic locations. Such locations have been called ‘hotspots’ by other authors, and many we have identified in this ST131 dataset were previously identified as region of high genetic turnover harboring defense systems (Hochhauser *et al*., 2023; Oliveira *et al*., 2017), indicating that diversification rates consistently and strongly depend on chromosomal location. This is consistent with the suggestion that the accumulation of defense systems into such ‘defense islands’ arises from the repeated insertion of MGEs at these locations (Rocha and Bikard, 2022).

Our results underscore the rapidity of genomic diversification: starting from the common ancestor of *E. coli* ST131 on the order of 100 years ago, large scale genome modifications have produced more than 500 structurally diverse regions that often harbour tens of different variations in a sample of ∼ 200 genomes. This diversity is likely to increase with the size of the dataset, given that a large fraction of the structural variants we observe are rare.

In contrast to the deletional bias described for bacterial genomes (Kuo and Ochman, 2009; Mira *et al*., 2001), we found that gains of genetic material were far more common than losses. This raises the question why bacterial genomes are stable in size and compact. These gains are mostly attributable to IS or prophage integrations that are likely mostly deleterious, especially when they disrupt functional genes. Even on the short time scale spanned by this dataset, two-thirds of insertions seem to have been removed by purifying selection. On the other hand, gene disruption or the introduction of new promoters by IS elements can also be adaptive, at least transiently (Sastre-Dominguez *et al*., 2024; Siguier *et al*., 2014). Such transient adaptive effects could contribute to an over-representation of gains on short time scales. In addition to purifying selection, the excess of insertions and their dominance on terminal branches (along with rearrangements) could also, in part, be due to recent bursts of IS activity in these genomes.

It is important to note that many of these gains of gene-disrupting IS elements would likely be counted as gene losses by traditional gene based pangenome tools (Marin *et al*., 2024; Tonkin-Hill *et al*., 2023). Similarly, the nucleotide based pangenome graph interprets local indel variation like SNPs, while a frame shift producing indel often results in “gene loss” in gene based pipelines (see *SI* fig. 22 and *SI* sect. 11). Correcting for this often requires integration with a graph-based approach (Horsfield *et al*., 2023; Tonkin-Hill *et al*., 2020).

Among the limitations of this work is the reliance on assembly accuracy. Assembly errors will introduce spurious structural diversity that is difficult to control for computationally. It is thus possible that some technical artefacts are interpreted as structural variation, leading to inflated rate estimates. By restricting the dataset to high quality reference sequences we sought to minimize this problem. The concordance of the structurally diverse junctions we identified with previously identified hotspots in broader collections of *E. coli* suggests spurious diversity is a minor factor. Another possible limitation is that our conclusions are based on *E. coli* ST131, a strain associated with hospital outbreaks and antimicrobial resistance. It is possible that ST131 represents a special case and that our findings do not generalise across all bacterial genomes or even all *E. coli* .

A more systematic analysis of the rapid diversification of “hotspots” would be a natural extension of this work. For example, one could quantify whether the observed diversity of these regions is compatible with nested gain/loss/translocation events on the phylogeny, or whether wholesale exchange with broader *E. coli* diversity is dominating over local diversification. Moreover, the analysis we conducted is well-suited to be extended to any recently-diverged microbial clade for which a reliable recombination-free phylogeny can be inferred. It will be interesting to see whether patterns of genome evolution differ qualitatively between different bacterial species.

We have shown that structural modifications of bacterial genomes happen at a rate comparable to that of individual nucleotide mutations anywhere in the genome. Extrapolating the dynamics described here for short timescales to larger and larger scales is a fundamental challenge for a quantitative understanding of the genotypic and phenotypic diversification of bacteria.

## REPRODUCIBILITY

The analysis contained in this paper can be fully reproduced with the pipeline available at https://github.com/mmolari/ecoliST131-structural-evo (v1.0).

## ACKNOWLEDGEMENTS

We are grateful for stimulating discussions with Erik van Nimwegen and Boris Shraiman and thank Eduardo Rocha and Zamin Iqbal for their helpful feedback on previous versions of this manuscript. This work was supported by the University of Basel (MM, RAN). LPS is a Sir Henry Wellcome Postdoctoral Fellow funded by Wellcome (grant 220422/Z/20/Z).

