## supplementary text and figures for "Evolutionary dynamics of genome structure and content among closely related bacteria"

Marco Molari,[1,2] Liam P. Shaw,[3,4] and Richard A. Neher[1,2,*]
[1]*Swiss Institute of Bioinformatics, Basel, Switzerland*
[2]*Biozentrum, University of Basel, Basel, Switzerland*
[3]*Department of Biology, University of Oxford, Oxford, UK*
[4]*Department of Biosciences, University of Durham, Durham, UK*

#### CONTENTS

I. Dataset creation and characterization 2
II. Pangenome graph construction 4
III. Core genome tree 6
IV. Quantifying the dataset diversity in terms of core and accessory genome 9
V. Core genome synteny 10
VI. Identification of loci (junctions) with accessory genome variation 13
VII. Functional characterization of accessory variation 14
VIII. Concordance of branch length and number of changes to genome structure 16
IX. Gene disruption by Insertion Sequences 17
X. Locating hotspots from Hochhauser et al. 18
XI. Comparison to gene-based pangenome analysis using panX 24
XII. Heterogeneity in rates of evolution of genome structure 25
References 26

### I. DATASET CREATION AND CHARACTERIZATION

#### A. Dataset Creation

The dataset was created by downloading all complete assemblies (complete chromosome and plasmids) from RefSeq (O'Leary *et al.*, 2016) on 26.09.2023 (n=3081). Of this we retain the subset that satisfies these two conditions:

- is assigned to ST131 by mlst v2.23.0 (Seemann, 2016) using the Achtman scheme (`ecoli_achtman_4`).
- has a mash (Ondov *et al.*, 2016) distance $< 0.008$ to the ST131 isolate `NC_013654.1`, used as reference in (Stoesser *et al.*, 2016).

This results in n=225 complete assemblies. We perform a quality check using BUSCO v5.5.0 (Manni *et al.*, 2021) and verify that no isolate has more than 2% of BUSCO genes missing or fragmented (average fraction of complete BUSCO genes: 99.96%).

After further quality control we decided to remove 3 more isolates from the dataset:

- in `NZ_CP093011.1` the core genome synteny is heavily fragmented. We suspect this might be due to assembly artifacts (isolate sequenced with Illumina MiSeq, coverage of 30x and assembled with SPAdes v. 3.11.1).
- isolates `NZ_CP124490.1` and `NC_013654.1` show signs of recombination in the core genome even after the recombination filtering procedure (see section III.A). This is detectable as a high number of homoplasies restricted exclusively to these two isolates.

The final dataset is composed of 222 isolates from all continents and spans a wide range of collection dates (see SI fig. S1). The large representation of sequences from Switzerland and Sweden are mainly due to these two studies: (Biggel *et al.*, 2023) and (Jaén-Luchoro *et al.*, 2023).

#### B. Dataset Characterization

We characterize the genomes in different ways, all shown in fig. 1 in the main text.

For typing of the *fimH*, *gyrA* and *parC* loci, we download the corresponding schemes from Enterobase (Zhou *et al.*, 2020) (respectively named `b4320`, `b2231` and `b3019`) and use blast to find and assign the best match. This is done by ranking hits between the genomes and the scheme sequences by number of exact nucleotide matches divided by the length of the scheme sequence. We only consider hits with $\geq 95\%$ identity, and in case of equality we take the longest scheme sequence. The vast majority of assigned types have 100% identity.

For characterising plasmids, we perform plasmid MLST using the IncF typing scheme downloaded from pMLST (Jolley *et al.*, 2018). As for the previous case we use blast to find the best match for each locus.

For resistance genes we use `abricate` v1.0.1 (Seemann, 2020) with CARD (Alcock *et al.*, 2023) database option.

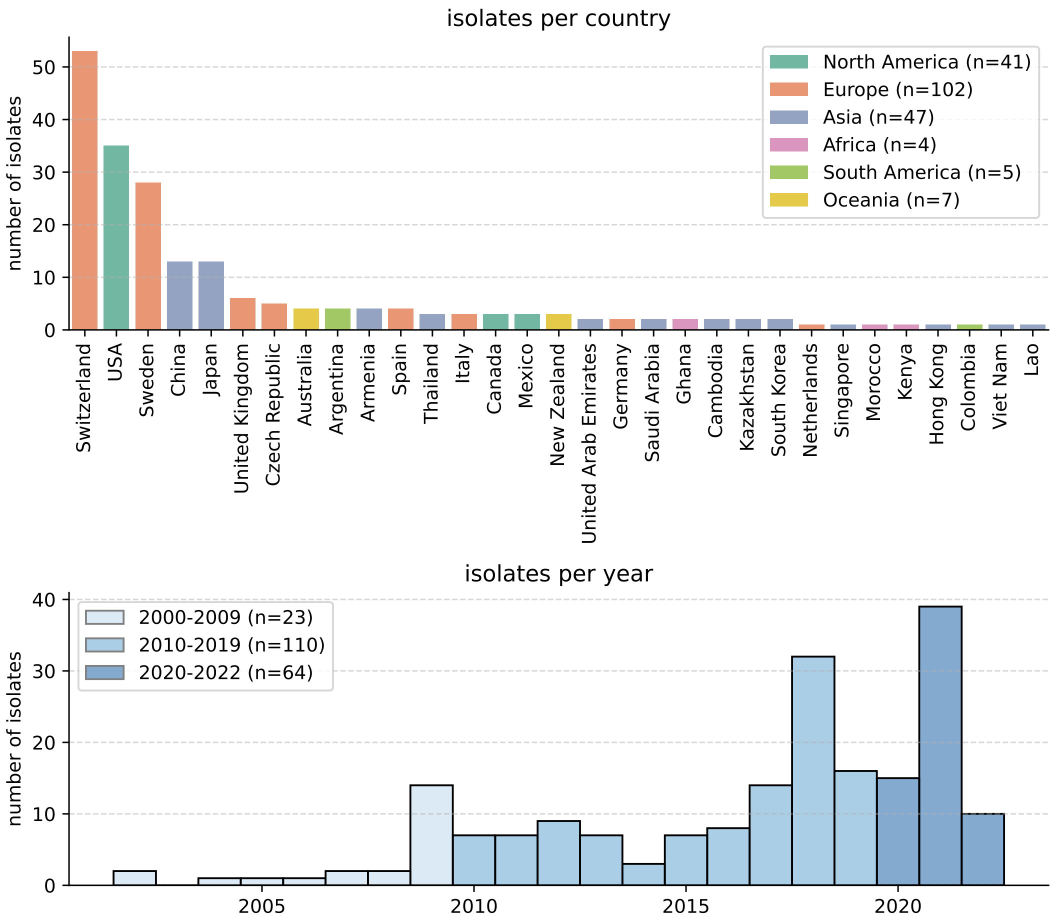


FIG. S1 **Top:** distribution of countries of origin of isolates in our dataset, for isolates for which this information is available. **Bottom:** distribution of isolation dates for our dataset, for isolates for which this information is available.

#### II. PANGENOME GRAPH CONSTRUCTION

We built a pangenome graph out of the 222 isolates in our dataset using PanGraph (Noll *et al.*, 2023). In this representation homologous parts of the sequence are stored in *blocks*, and genomes are represented as *paths* through these blocks, see fig. S2. We use PanGraph version 0.7.3, with options:

- `--circular`: to account for the circular nature of bacterial chromosomes.
- `-k minimap2 -s 20`: we use *minimap2* (Li, 2018) as alignment kernel, with sensitivity option `asm20`, which is suitable for align genomes with up to 5% divergence (Noll *et al.*, 2023), well above the average divergence of our dataset (∼0.01%).
- `--alpha 100`: block fragmentation cost. Whenever an incomplete match generates flanking blocks in the merging, for the match to be accepted its length should be at least 100 bp per flanking block created. This avoid excessive fragmentation.
- `--beta 5`: divergence threshold for block merging. Blocks are merged if they are separated by less than 5% divergence. This divergence threshold is chosen to be consistent with the sensitivity of the alignment kernel.
- `--len 100`: we use the default value for the block threshold length. Matches shorter than this length are not merged into new blocks.

In fig S3 we display the distribution of block frequency (i.e. number of isolates where the block is present) and length. The pangenome graph is composed of 4422 blocks, and has total pangenome size of 7.99 Mbp. Of these, 631 are single-copy core blocks (i.e. present exactly once in every isolate) and these constitute roughly half of the pangenome in terms of size, for a total of 3.59 Mbp. If we extend this sum to soft-core blocks (present in $> 95\%$ of isolates) we get a total size for the soft-core genome of 4.16 Mbp. The large proportion of core genome relative to the full pangenome can be attributed to the high relatedness between these isolates. The graphs also contains 1303 duplicated blocks. These are blocks that appear more than once in at least one isolate. These are generally short blocks, and in total they add up to around 0.8 Mbp.

Blocks have a median length of around 600 bp. However if weighted by length, most of the sequence in the graph is included in blocks of size of almost 10 kbp. Most of these long blocks are core blocks, and are present in all isolates.

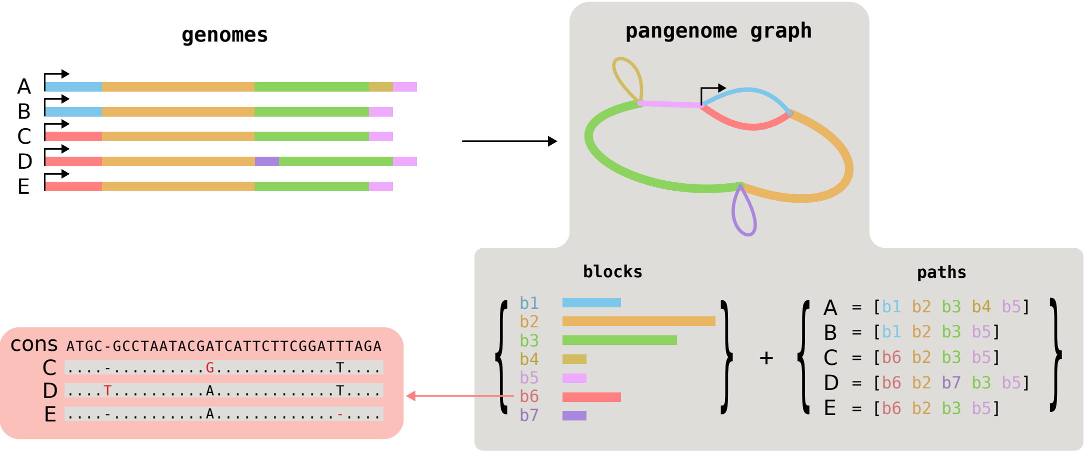


FIG. S2 **Schematic depiction of the pangenome graph created by *PanGraph*.** Given a set of genomes, PanGraph can build a *pangenome graph* representation for the genomes. This representation consists of *blocks*, encoding multiple sequence alignments of homologous regions, and *paths*, encoding genomes as sequences of blocks. Each block is defined by a consensus sequence, and a set of variations over the consensus for each occurrence of the block.

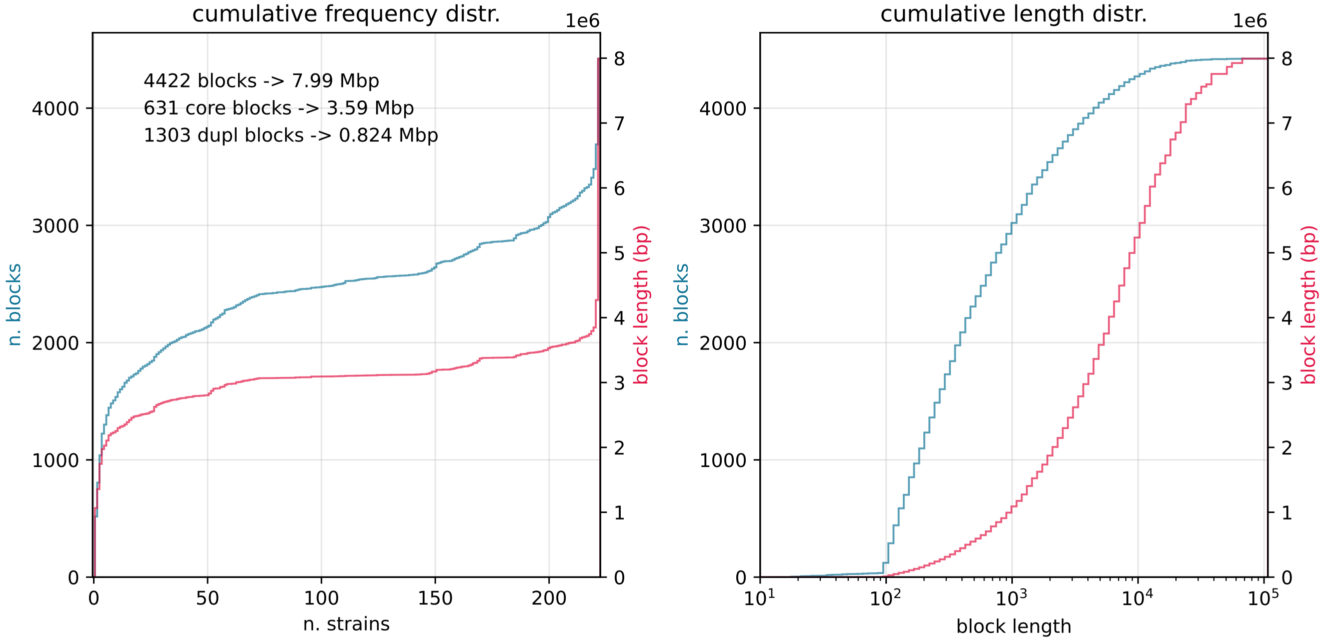


FIG. S3 **Cumulative distributions of block frequencies (left, i.e. number of isolates in which the block is present) and lengths (right) for the pangenome graph.** Distributions are reported both in number of blocks (red) and weighted by block length (blue). Additionally, on the left panel we report the total number of blocks and total pangenome size, toghether with the number and total size of core and duplicated blocks.

#### III. CORE GENOME TREE

##### A. Core genome alignment and recombination filter

In order to build the core genome graph from our isolates, we concatenate all of the core block alignments in a single alignment having total length of 3.59 Mbp. The blocks are ordered in the alignment using the order of isolate `NZ_CP096110.1`, which has the consensus synteny.

This alignment contains 21939 polymorphic sites (counting only positions without gaps). These sites can be used to build a phylogenetic tree. Homologous recombination, both within our dataset and with other microbes, can however bias the reconstructed tree, both in topology and in branch length. In order for this tree to be as close as possible to the cell-division history of the population we need to remove these recombined regions. Since the isolates in our datasets are highly similar, it is likely that homologous recombination with other microbes resulted in densely mutated regions (Sakoparnig *et al.*, 2021). We therefore filter the alignment using the following simple procedure:

1. We remove from the alignment any position containing a gap or an ambiguous nucleotide in any isolate.

2. We then consider isolates separately, and for each position that is mutated w.r.t. the consensus we count the number of other mutated sites within 1 kbp.

3. If the number of such sites is larger than 3 on a single isolate, we remove all positions within 1 kbp from each of the mutated sites from the whole alignment. This removes regions in which at least one isolate has many clustered mutations.

The resulting filtered alignment has size of 2.43 Mbp. The number of polymorphic sites in the filtered alignment decreases to 5979. The effect of this filtering procedure is displayed in fig. S4. In particular regions with high SNPs density, where most of the mutations are contained, are removed, leaving regions that have the background mutation density of roughly one polymorphic site per kbp over our dataset. This filter renders the mutation density homogeneous along the alignment (cf. fig. S4 bottom). On this filtered alignment, the average pairwise divergence of our isolates is 74 mutations per Mbp, mainly attributable to cross-clade comparisons.

We build a core genome tree from this refined alignment using *FastTree* (v2.1.11) (Price *et al.*, 2010), and then polish the tree using TreeTime (Sagulenko *et al.*, 2018) (v0.9.4).

We also verified that the tree obtained with our simple recombination filtering procedure is very similar, both in branch length and topology, to the one obtained by filtering recombination with Gubbins (Croucher *et al.*, 2015) (v3.3.1), which uses a more complex procedure.

##### B. Testing the clonality of the tree using homoplasies

The procedure we used allows us to remove recombination with distantly related microbes, but is not able to detect homologous recombination within our sample. To verify that our filtered alignment is largely compatible with clonal evolution we check for the amount of homoplasies in the alignment. The reason is that horizontally transferred mutations would be visible as homoplasies under the false hypothesis of clonal evolution in our inferred tree. If our recombination filter is effective in removing also possible within-sample recombination, then it must remove most of the homoplasies from the alignment.

To verify this we consider the core genome alignment before and after the filtering procedure. For each of these cases we infer a core genome tree using the procedure specified above, and then perform ancestral sequence reconstruction using TreeTime (Sagulenko *et al.*, 2018). We infer two separate trees in order for them to be maximally compatible with the alignment we are considering, minimizing the amount of homoplasies.

To meaningfully define homoplasies we only consider mutations that are:

- bi-allelic, i.e. present in exactly two different states in the alignment.

- non-singleton, i.e. present in at least two isolates.

These are mutations that can be meaningfully incompatible with a single event on our tree. We define homoplasic mutations as the ones from this category that are inferred to occur more than once on our tree.

For this inference we consider the core genome alignment restricted to variable sites that do not contain gaps. There are 21939 such sites in the pre-filter alignment and 5979 in the post-filter alignment. This different length is expected given the action of the filter, which removes highly-mutated regions. In SI fig. S5 we show the fraction of bi-allelic

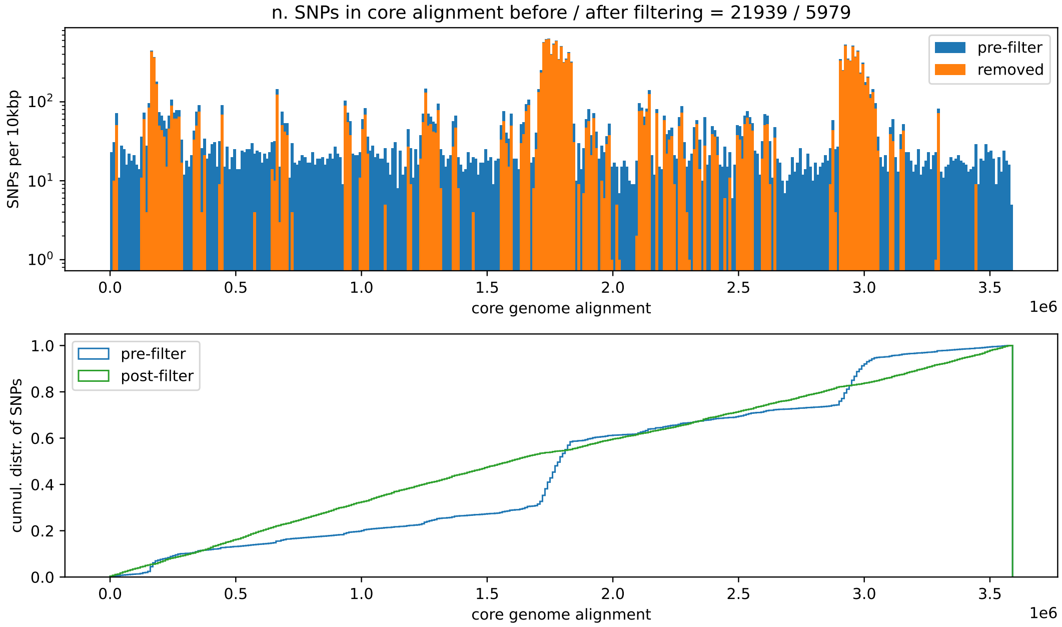


FIG. S4 **Effect of the recombination filter on the core genome alignment.** The filter removes regions of high SNPs density. **Top**: total number of SNPs in 10 kbp bins on the core genome alignment (blue) and amount of SNPs removed by the filter (orange). **Bottom**: cumulative distribution of SNPs along the original core genome alignment before (blue) and after (green) filtering.

and non-signleton mutations in the restricted alignment, stratified by whether they are homoplasies. The average fraction of such mutations is larger in the pre-filter alignment (62%) than in the post-filter alignment (20%). The vast majority of the remaining mutations in the two restricted alignments are still bi-allelic but the minority allele only occur in a single isolate (singleton mutation).

Importantly, after the recombination filter the fraction of homoplasic mutations decreases from 35% in the pre-filter alignment to only less than 3% (n=33) in the post-filter alignment. For these few remaining homoplasies we verified that their pattern on the tree is sparse and that they cannot be attributed to few highly-recombined isolates.

Given this observation we are confident that our filter is effective in removing most of the recombination signal, and that the inferred core genome tree is a good approximation of the vertical evolution history of our isolates.

##### C. Syntenic core genome blocks

To analyze core genome synteny, we constructed reduced paths containing only core blocks longer than 500bp by skipping accessory blocks or short blocks. This additional length constraint only removed few short blocks (n=95) and was chosen to ensure that all blocks considered are well above the length cutoff parameter of PanGraph (set to 100 bp). We then merged any two core-blocks that always appear in the same order and with the same strandedness. We thereby obtained 32 maximal blocks that are syntenic across all isolates in the dataset, and whose end-points correspond to positions where the core genome order differs between isolates in our dataset.

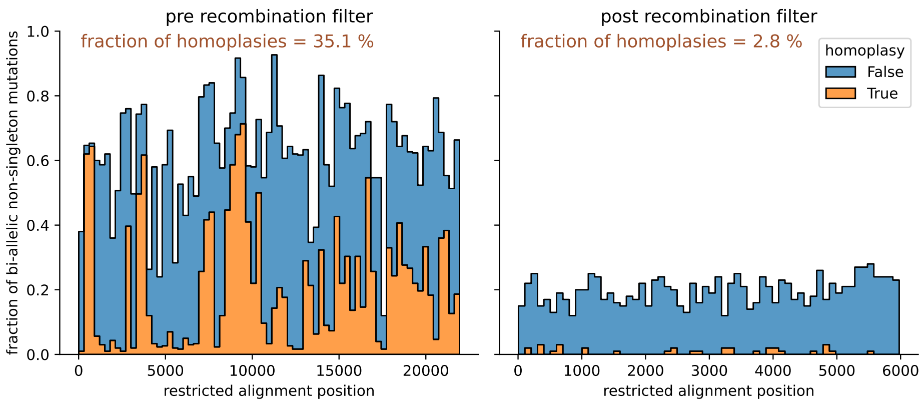


FIG. S5 **Fraction of bi-allelic and non-singleton mutations in the restricted core genome alignment.** Mutations are stratified by whether they are homoplasies (stacked histogram). The restricted alignment only contains non-consensus sites. **Left**: pre recombination filter. **Right**: post recombination filter.

#### IV. QUANTIFYING THE DATASET DIVERSITY IN TERMS OF CORE AND ACCESSORY GENOME

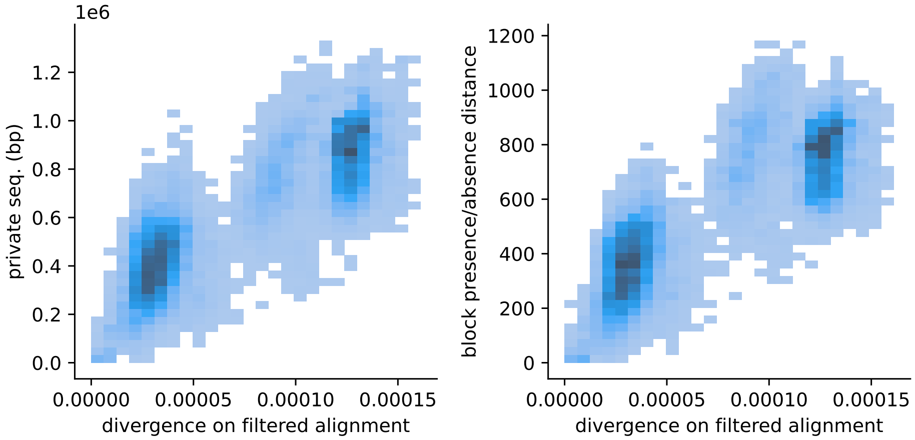


FIG. S6 **Quantification of the diversity of the dataset with three different distance measures.** For every pair of isolates we quantify: 1) the average sequence divergence over the filtered core genome alignment, i.e. the fraction of SNPs over the total alignment length, excluding sites with gaps. 2) the amount of private sequence, i.e. the total length of blocks present in either one of the isolates from the pair but not the other. 3) the block presence/absence distance, i.e. the number of blocks present in the path of either one of the isolates but not the other. For all pairs in the dataset we display the joint distribution of average sequence divergence and amount of private sequence **(left)** and the joint distribution of average sequence divergence and block presence/absence distance **(right)**. While being extremely similar on the core genome (avg. divergence $7.4 \times 10^{-5}$) isolates in our datasets have large differences in accessory sequence (on average 610 kbp of private sequence, and 545 blocks of difference).

#### V. CORE GENOME SYNTENY

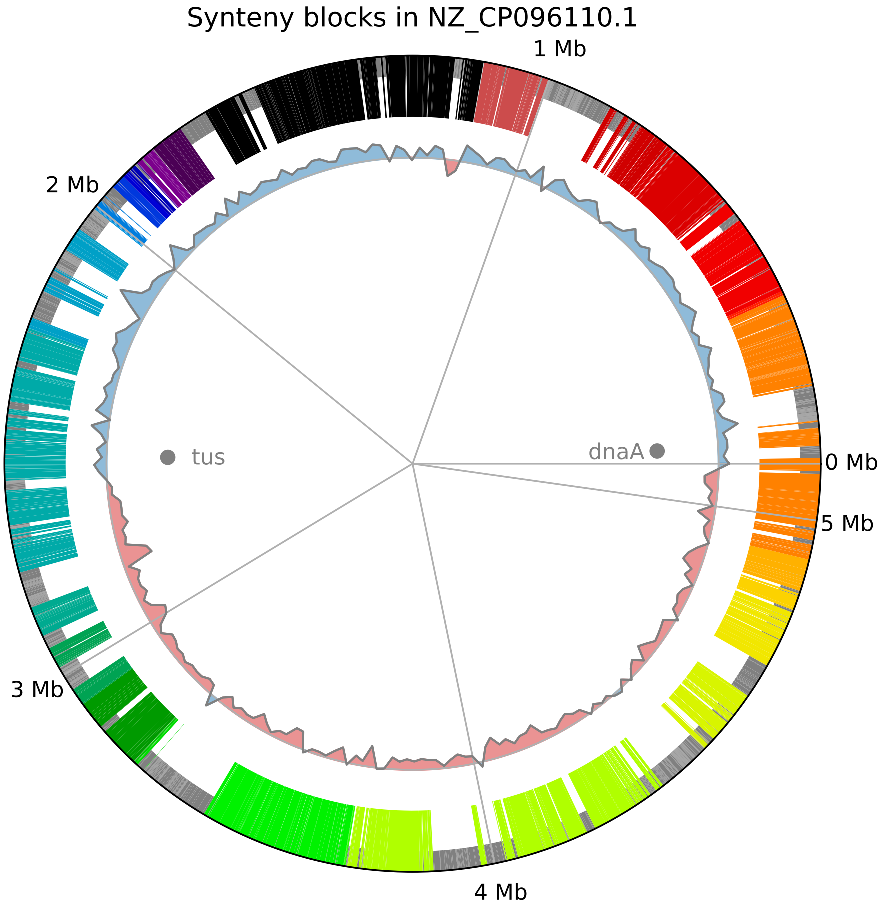


FIG. S7 **Projection of synteny blocks on reference sequence** `NZ_CP096110.1`**.** Accessory blocks are represented as thin grey boxes, while core blocks are thicker and colored according to the minimal synteny block they belong to, as in fig. 2 in the main text. The inner circle measures the GC-skew in bins of 20 kbp, indicating the origin and terminus of replication. We also mark the positions of the *dnaA* and *tus* genes, close to the origin and terminus of replication.

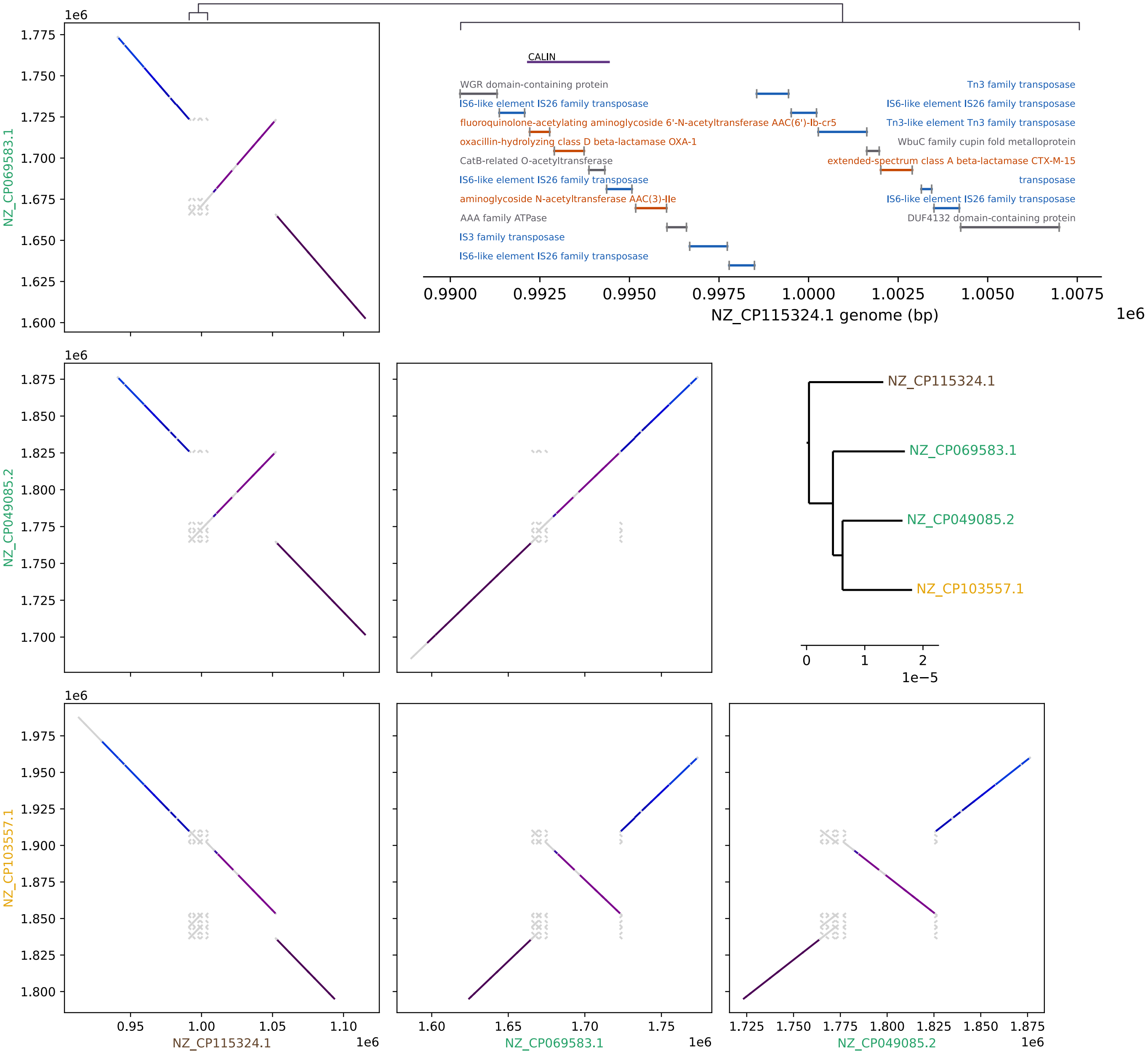


FIG. S8 **Comparative analysis of the inverted region in isolates** `NZ_CP069583.1` **and** `NZ_CP049085.2` (green). These two isolates present an inversion that is absent in the third member of the clade `NZ_CP103557.1` (yellow). This might be due to either errors in phylogenetic reconstruction, or to the reversion of the inversion. Comparison of the structure of this region between the three isolate and the closest outgroup sequence `NZ_CP103557.1` (brown) reveals the appearance of an extra copy of a duplicated region flanking the inversion in `NZ_CP103557.1` (yellow), suggestive of a reversion of the inversion. This region contains several ISs, a CALIN annotation and several anti-microbial resistance genes. **Top right**: annotations of the region around 1Mb in the outgroup isolate `NZ_CP103557.1`. We indicate in blue genes annotated as ISs and transposases, in orange genes related to AMR, and in purple we indicate the region annotated as CALIN by IntegronFinder. **Center right**: Clade containing the isolates of interes. The two isolates in green contain the inversion. **Bottom left**: dotplot obtained using the pangenome graph around the inverted region. For each pari of isolates, shared blocks are displayed as lines that connect start/end point of the block on each genome. Core-blocks are colored as in fig. S7, while accessory blocks are grey.

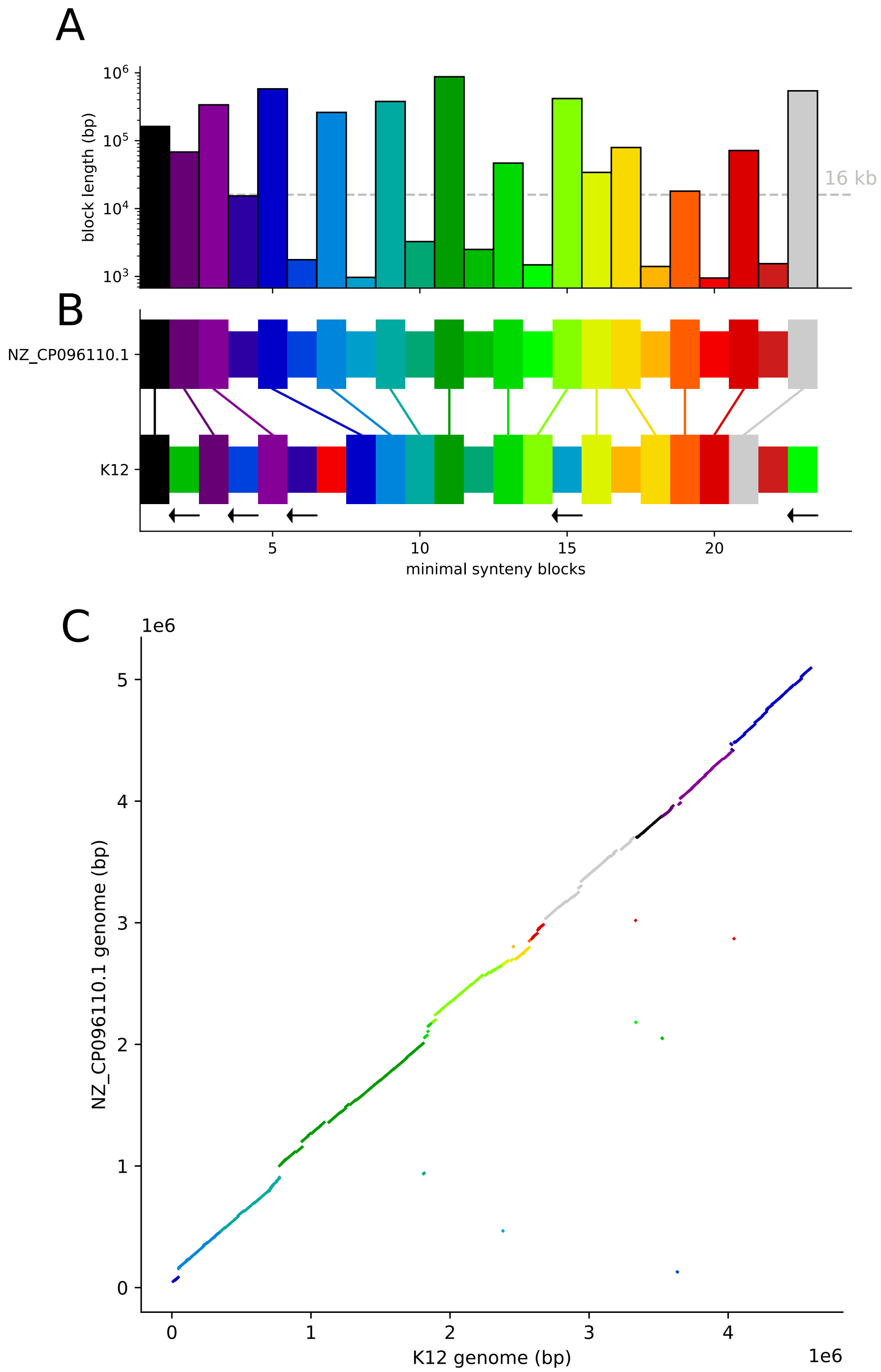


FIG. S9 **Comparison of core genome synteny for *E. coli* ST131 isolate `NZ_CP096110.1` and *E. coli* K-12 substrain MG1655 (acc. number `CP014348.1`).** We build a pangenome graph for the two isolates. The total amount of core-sequence for the pair (single-copy shared sequence) is 3.9 Mbp, with an average sequence divergence of 2.7%, roughly 200 times higher than the average for the ST131 dataset. As done for the ST131 dataset, we filter out from paths all non-core blocks and blocks shorter than 500 bp and then merge transitive edge to obtain minimal synteny blocks. Core genome synteny is highly consistent. The two core genomes differ only by the movement of 9 "short" blocks. **A**: size of minimal synteny block for the comparison between the two genomes. The horizontal dashed lines separates "short" and "long" blocks. **B**: comparison between order of minimal synteny blocks in the two genomes. Inversion is indicated with a black arrow below the block. The block height indicates whether the block is "long" or "short". Notice how the order of all of the "long" blocks is conserved. **C**: dotplot for the core blocks shared between the two genomes. The color of each core block represents the minimal synteny block it belongs to.

#### VI. IDENTIFICATION OF LOCI (JUNCTIONS) WITH ACCESSORY GENOME VARIATION

To systematically quantify variations in the accessory genome we use the blocks of the core genome as a frame of reference. We define "core-edges" as follows:

- as for the synteny analysis, we remove from each path all non-core blocks and core-blocks shorter than 500bp.
- on these stripped-down core-paths every pair of consecutive blocks $A$ and $B$, together with their strandedness ($+$ or $-$), defines a core-edge, e.g. $(A+, B+)$.

Core-edges are equivalent under reverse-complementation (i.e. $(A+, B+) = (B-, A-)$). Each equivalence class defines a conserved context in which we can compare genomic changes. Due to synteny breaks, any given core-edge is not necessarily present in every isolate. We can split core-edges in three categories, see fig. S10:

- *backbone edges* (n=504): present in all 222 isolates.
- *common edges* (n=32): present in all isolates apart from a small minority that have a different synteny pattern.
- *rare edges* (n=45): present only in a minority of isolates in correspondence of a synteny break.

Notice how out of 536 core blocks ($> 500$ bp) in the graph, 504 pairs always follow one another in the same order, but with different accessory content in between.

To study the accessory variation for each core-edge we extract from the genome of each isolate all the sequence from one core block to the other, including the flanking core blocks themselves. We then re run PanGraph on these sequences with the same options used to produce the main graph, with the exception of the `--circular` flag. This generates smaller *junction graphs*, see fig. fig 3A in the main text. For each of these graphs we count the number of different distinct paths. Junction graphs with a single possible path are defined as *transitive* edges/junctions. Other than in the trivial case of edges present in only one isolate, this can also happen for example when two core blocks are separated by an element that is duplicated somewhere else in the genome (and thus is classified as accessory sequence) but this element is fixed in the population in this particular position. Since these junctions do not effectively contain diversity across the dataset we exclude them from the next steps of the analysis. In fig. S10 we display the frequency distribution of core-edges in our dataset.

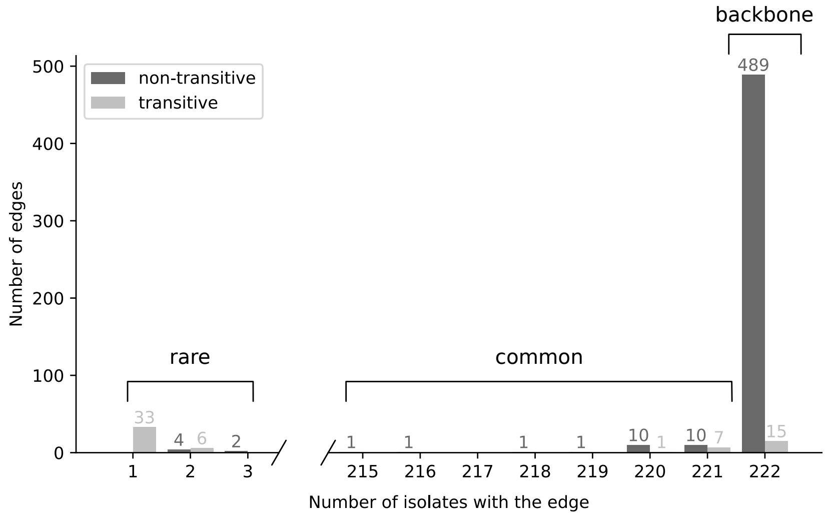


FIG. S10 **Frequency distribution of core-edges in our dataset.** Color indicates whether the edge is transitive or not. Most edges are *backbone*, i.e. present in all isolates.

#### VII. FUNCTIONAL CHARACTERIZATION OF ACCESSORY VARIATION

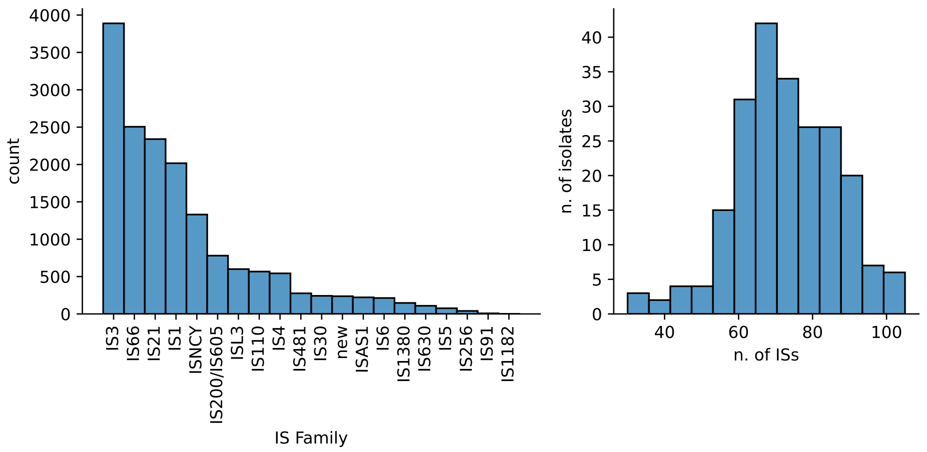


FIG. S11 **Distribution of Insertion Sequence annotations in our dataset**. ISs are detected by *ISEScan* (Xie and Tang, 2017). **Left**: distribution of IS families. **Right**: distribution of total number of IS per chromosome.

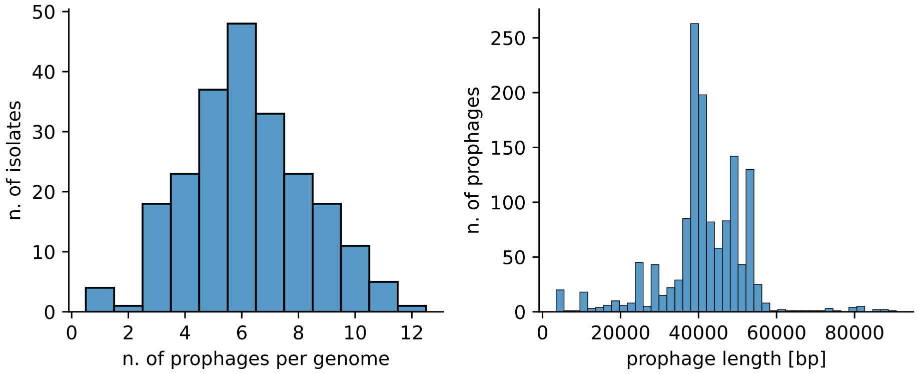


FIG. S12 **Distribution of prophage annotations**. Prophage detected by *GeNomad* (Camargo *et al.*, 2023). **Left**: distribution of number of prophages per chromosome in our dataset. **Right**: distribution of prophage length.

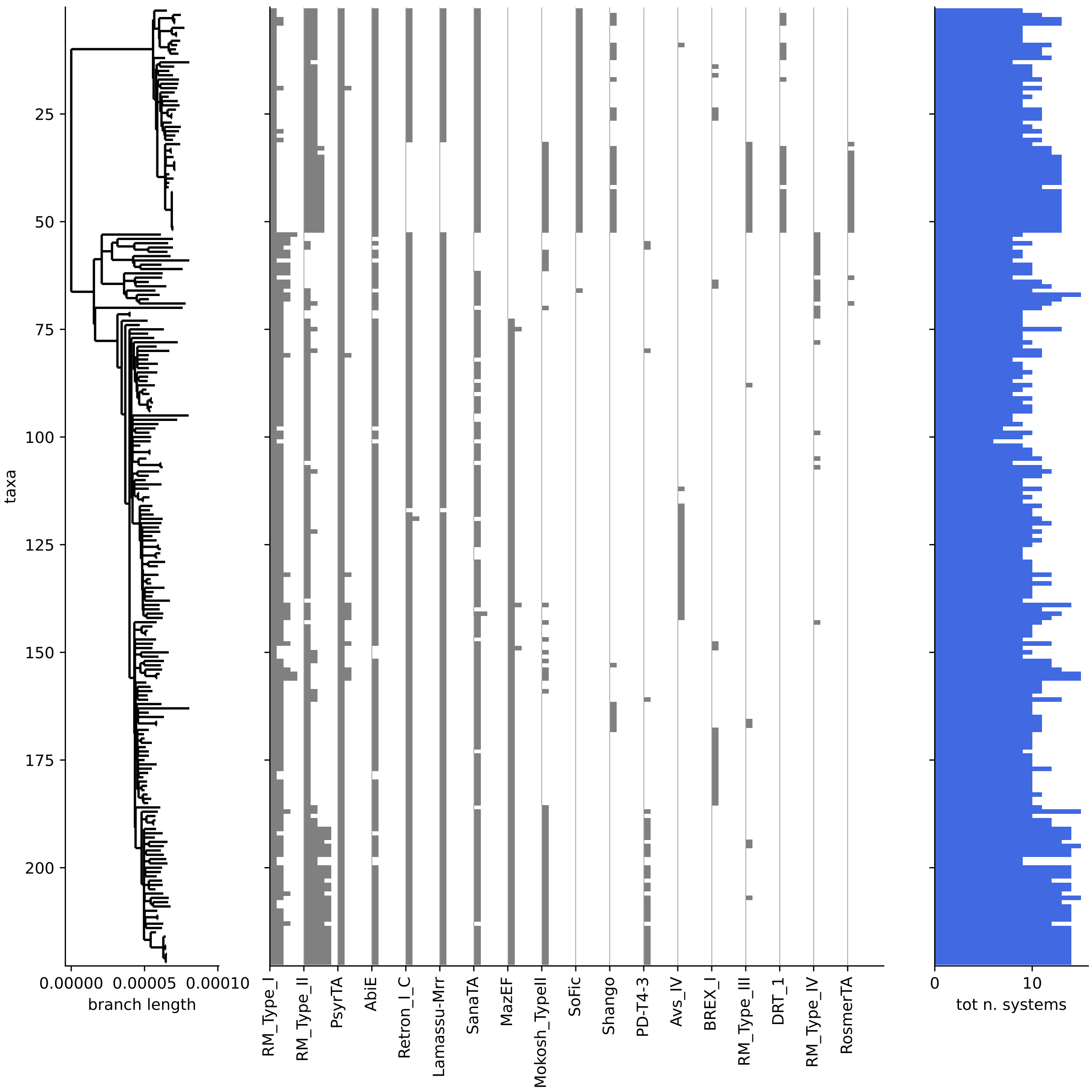


FIG. S13 **Distribution of defense systems in our dataset**. Defense systems detected by *DefenseFinder* (Tesson *et al.*, 2022). We detect a total of 2427 systems, with an average of 11 systems per isolate. These systems tend to localize in few long and complex junctions (41 junctions, 7% of all junctions). **Left**: core genome tree. **Center**: number of defense systems per isolate for the 18 most common defense systems, as identified by *DefenseFinder*. **Right**: total number of defense systems per isolate.

#### VIII. CONCORDANCE OF BRANCH LENGTH AND NUMBER OF CHANGES TO GENOME STRUCTURE

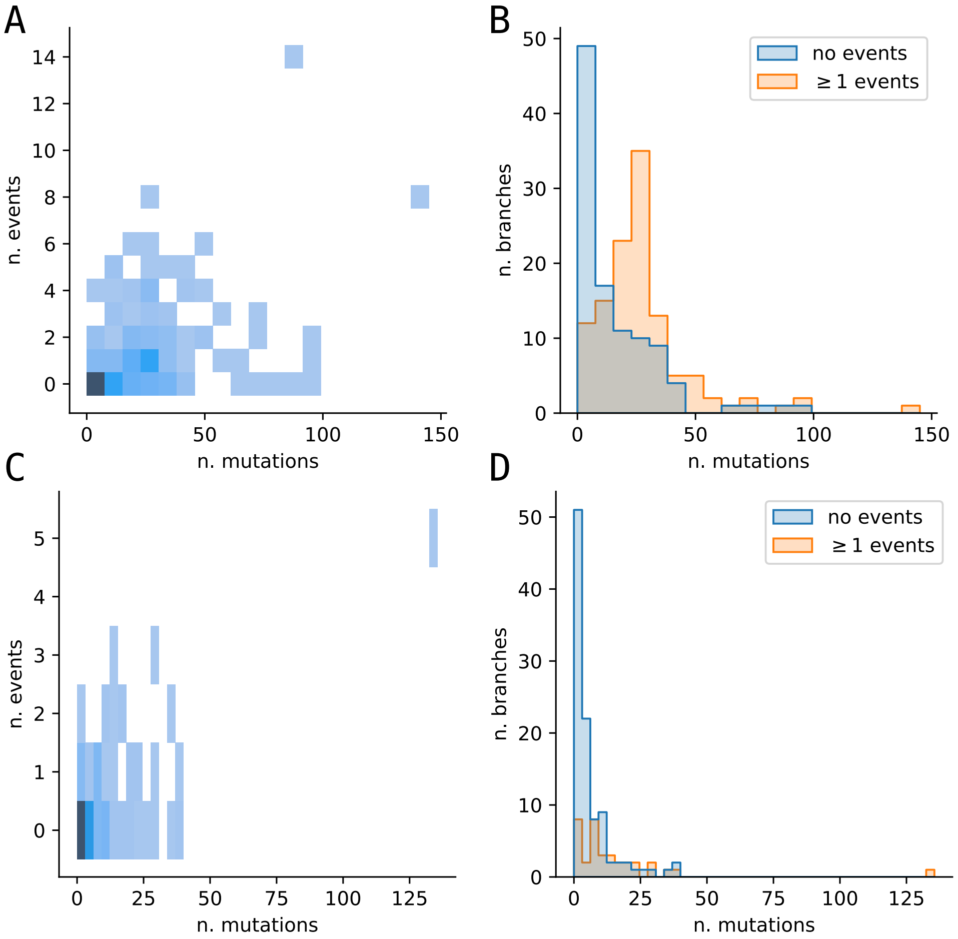


FIG. S14 **Correlation between branch length and number of gain and loss events associated to the branch.** On the left we display the joint distribution between the number of mutations associated to a branch (i.e. branch length times total filtered core-alignment size) and the number of events assigned to the branch for *terminal* (**A**) and *internal* (**C**) branches. On the right we display the distribution of number of mutations associated to a branch for *internal* (**B**) and *terminal* (**D**) branches, stratified by whether at least one event is assigned to the branch.

#### IX. GENE DISRUPTION BY INSERTION SEQUENCES

To verify the major role played by ISs in gain events, and to quantify the impact on gene disruption, we perform the following analysis:

- out of all backbone binary junctions, we select the ones whose accessory region contains at least one IS annotation.
- we then verify that these IS annotations are compatible with the set of distinct paths identified by pangraph. We can divide the isolates according to whether the accessory region in their path is empty or not, and whether they are associated or not to an IS annotation. Complete consistency requires that these two subdivisions coincide, with non-empty paths being the IS-associated ones.
- for these consistent junctions, we consider the isolates with an empty path. We check whether a gene annotation is present on the breakpoint defined by the position at which in the non-empty paths an IS was inserted.
- For those isolates in which the IS disrupts a gene, we verify whether the disrupted gene has the same name across all isolates.

Results for each of these steps are displayed in fig. S15. Interestingly, the fraction of IS-associated breakpoints that disrupt a gene is 71% (188/265), while insertions in non-coding regions are found in 29% of cases. Even though this is a considerable fraction, it is less than expected by chance. Across our dataset non-coding regions account on average for 11.6% of the genome. The fraction of observed non-coding insertions is roughly a third of what expected by chance. In the hypothesis that most of non-coding insertions are neutral, while insertion in coding regions are mostly deleterious, this suggests that roughly 2/3 of IS insertions in coding regions are removed by purifying selection and not observed.

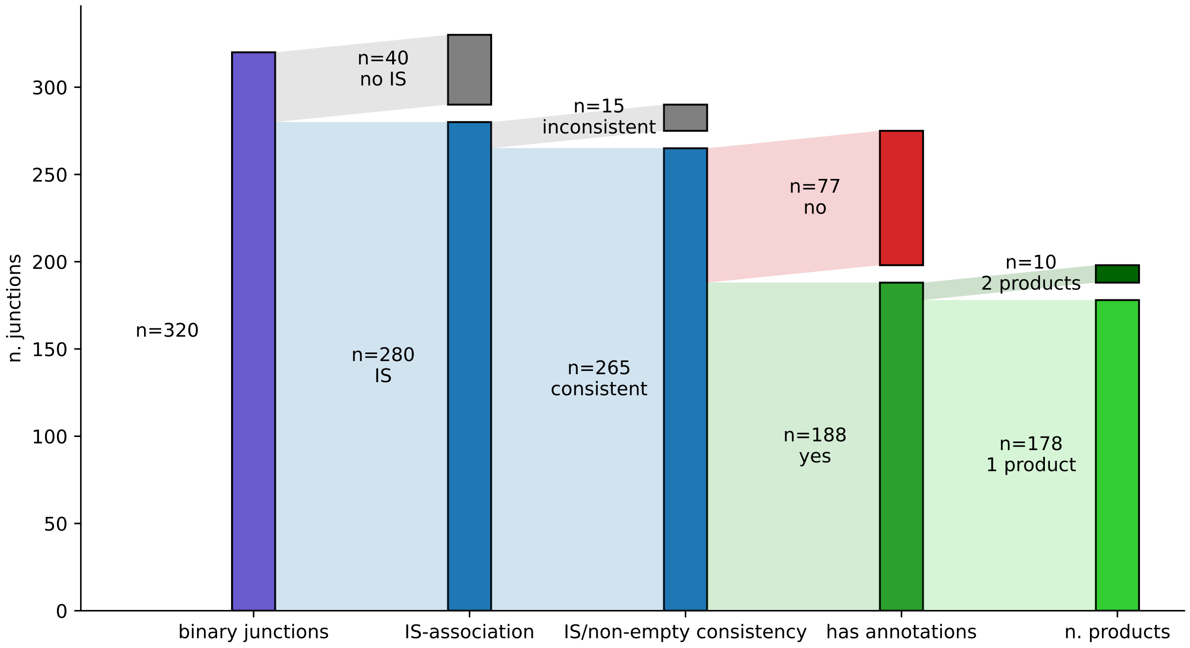


FIG. S15 **Association between insertion sequence gains and gene disruption.** Following the procedure indicated in the text, we quantify out of all of the backbone binary junctions, how many are associated to IS, in how many this association annotations are consistent with PanGraph paths, in how many cases the IS disrupts a gene, and in how many cases the disrupted gene is the same on all isolates.

#### X. LOCATING HOTSPOTS FROM HOCHHAUSER ET AL.

In (Hochhauser *et al.*, 2023) the authors study locations of preferential integrations of defense systems in a collection of > 1300 *E. coli* genomes. These integration sites are mapped to 41 *hotspots* in the genome of *E. coli* K-12 substrain MG1655, identified by the pair of closest flanking core genes. These spots are not present in every isolate or they are sometimes empty, but when present and occupied they often carry defense systems.

We can use the identity of the flanking core gene to locate these hotspots in our dataset, and check whether they correspond to some of the complex junctions that we identified. To do so we start by mapping the nucleotide sequence of the flanking core genes on the genome of isolate `NZ_CP096110.1` using minimap2. We only consider mappings with more than 90% of the expected sequence preset. We are able to identify 70/82 flanking core genes, with a total of 30/41 hotspots identified, see fig. S16. On this reference isolate only few of these hotspots are occupied. Moreover we verified that the located flanking core genes are either found on core-blocks (46/70) or on accessory blocks with frequency > 97%.

We then proceeded to assign these regions to junctions. We assign a hotspot to a junction if the region spanning the two flanking genes crosses the accessory part of a junction.

Out of the 30 locatable hotspots:

- 2 spots (n. 23 and 30) lay completely in a core-block and are thus not associated to any accessory junction. They correspond to empty spots.
- 1 spot (n. 6) is associated to a non-backbone junction, and in some isolates the synteny between the two flanking core genes is broken.
- two spots (n. 37 and 14) span two junctions.
- the remaining 25 spots are uniquely associated to a junction.

The association between hotspots and junctions is displayed in fig. S17. Note that with a few exceptions, most of the complex junctions that we identify corresponds to hotspots identified by Hochhauser et al. There are two junctions that are associated to two (n. 14 and 15) and three (n. 27, 28 and 29) hotspots. In the first case this is because the

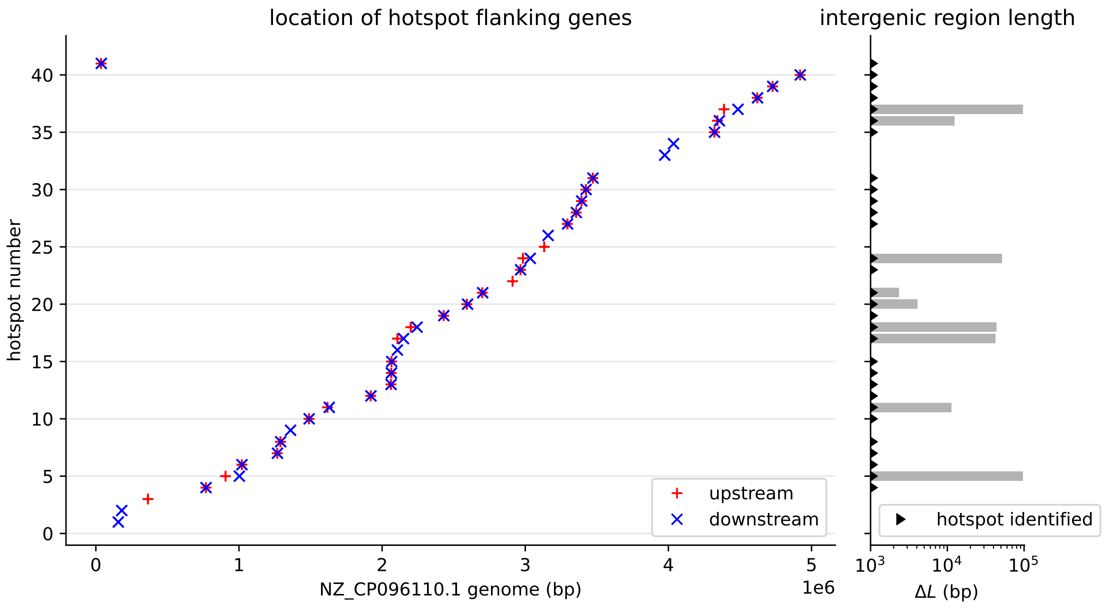


FIG. S16 **Location of hotspots on the genome of isolate `NZ_CP096110.1`.** **left**: location of upstream and downstream hotspot flanking genes on the genome. Note that for some hotspots some flanking genes could not be located. **right**: length of intergenic region for hotspots with more than 1kbp of sequence. This indicates which hotspots are occupied by accessory sequence on this genome. Hotspots that were identified (both upstream and downstream genes present) are marked with a black triangle.

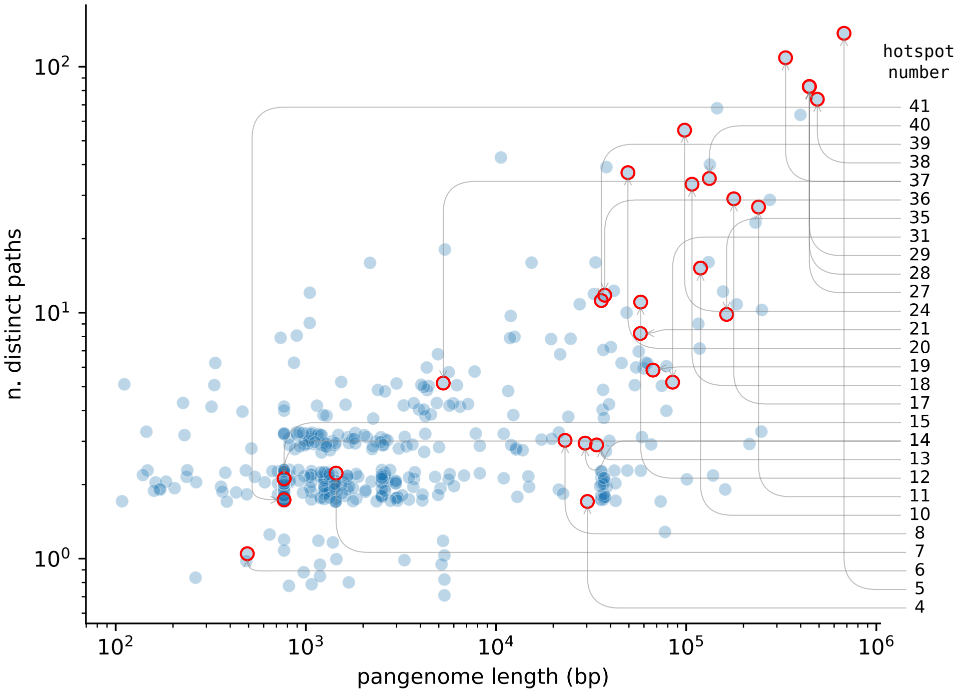


FIG. S17 **Association between junctions in our pangenome graph and hotspots identified by Hochhauser et al.** Junctions overlapping with a hotspot are marked with a red dot. Arrows indicate the connection between these junction and the hotspots, numbered as in (Hochhauser *et al.*, 2023).

flanking genes are the same, and in the second because the flanking genes are not core and are all found within a large accessory junction.

To illustrate the level of structural diversity that is found in complex junctions we picked two examples, corresponding to the junctions containing hotspots 18 and 11, and created two visualizations for each. Each line in the visualization represents the junction on a single isolate, ordered according to the phylogeny on the left. Black and red genes indicate the flanking genes used in (Hochhauser *et al.*, 2023) to identify the hotspot, and orange genes are tRNA genes, known for being targets of many integrase enzymes (Bellanger *et al.*, 2014; Williams, 2002). On top of this representation we highlight either the presence of different MGEs and defense systems (fig. S18 and S20) or the block structure defined by PanGraph (fig. S19 and S21).

Hotspot 18 (see figs. S18 and S19) presents a clear split between clade A and the rest of the dataset. In the former the hotspot is empty, except for a single isolate with a prophage integration. In clade B and C instead a different prophage is integrated. This prophage sequence presents nested diversity, mostly attributable to the movement of insertion sequences. In hotspot 11 (see figs. S20 and S21) we find instead the integration of many defense systems in clade A, and of few different prophages in clade B and C. Also in this case, the movement of insertion sequences adds more small-scale structural diversity.

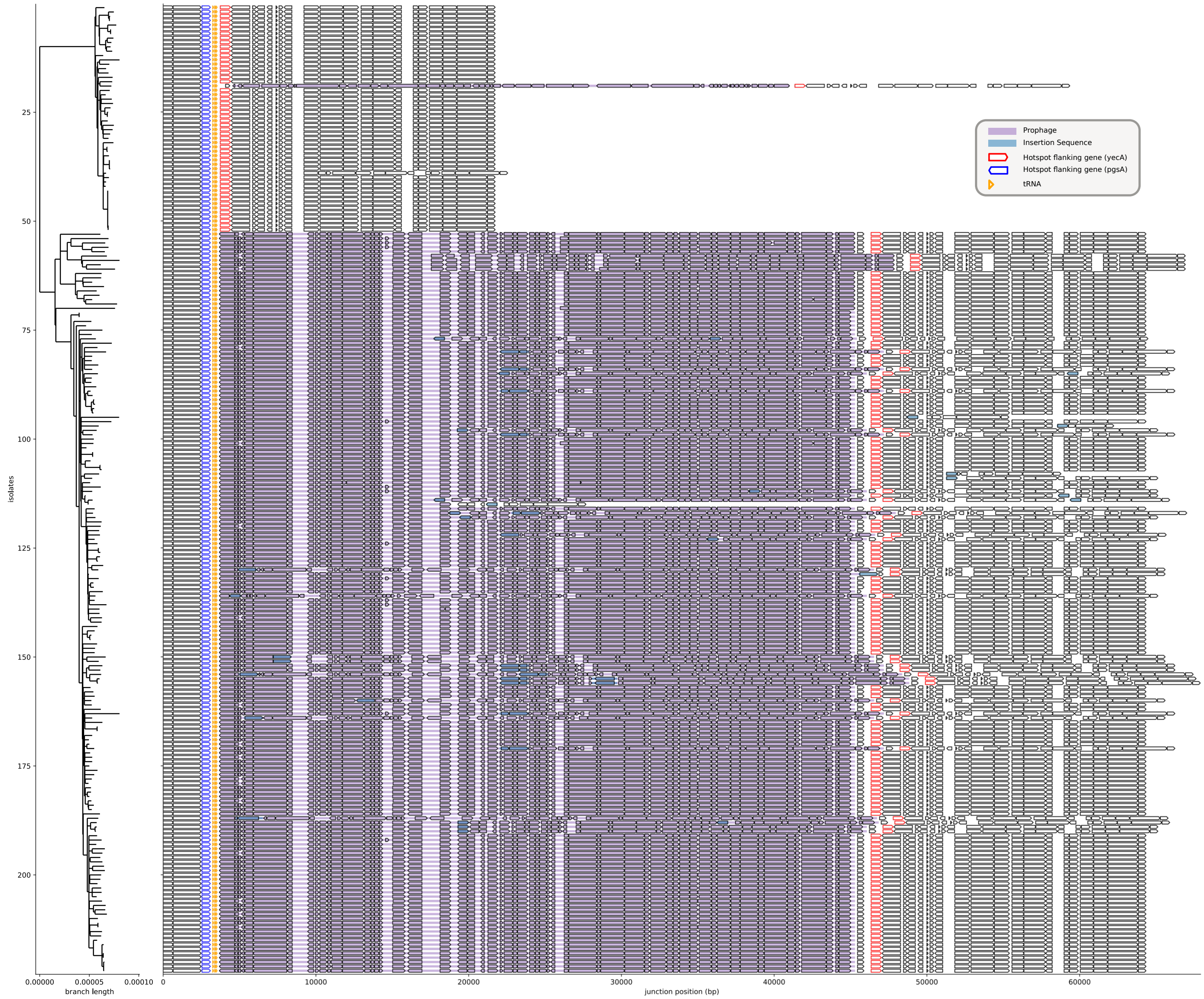


FIG. S18 **Example of functional content of complex junction containing hotspot 18**. Linear representation for the genomic region contained in a complex junction featuring 33 different distinct paths and a total pangenome length of 107 kbp. This junction contains the region named *hotspot 18* in (Hochhauser *et al.*, 2023), identifiable by its flanking genes (blue and red genes). Orange annotations are tRNA genes, while colored regions highlight annotations associated to prophages (purple) and insertion sequences (blue).

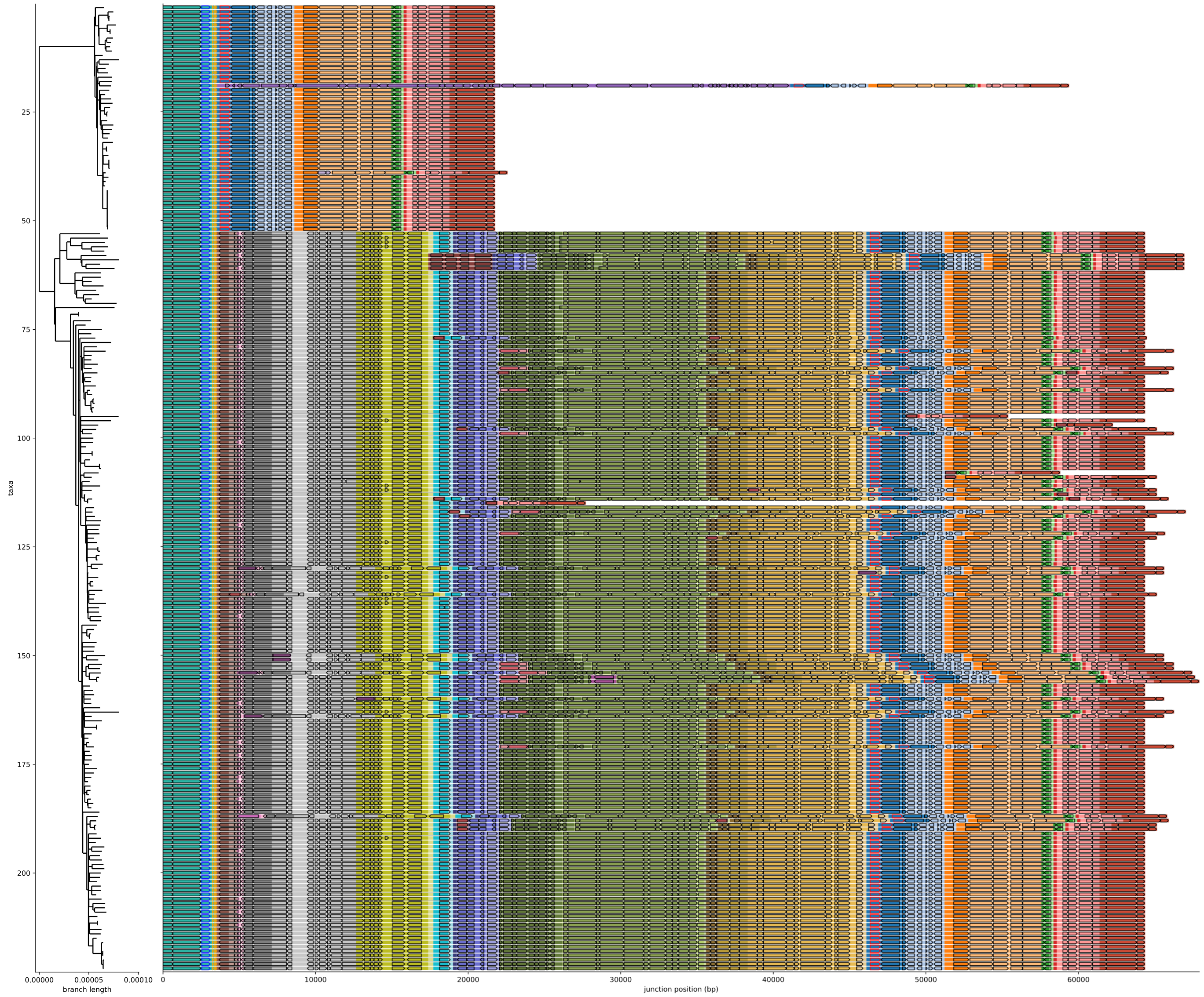


FIG. S19 **Example of structural diversity of a complex junction containing hotspot 18**. The figure contains the same junction depicted in fig. S18, but with colors indicating regions belonging to different blocks in the pagenome graph representation. The first (teal) and last (red) block correspond to the flanking core blocks of the junction.

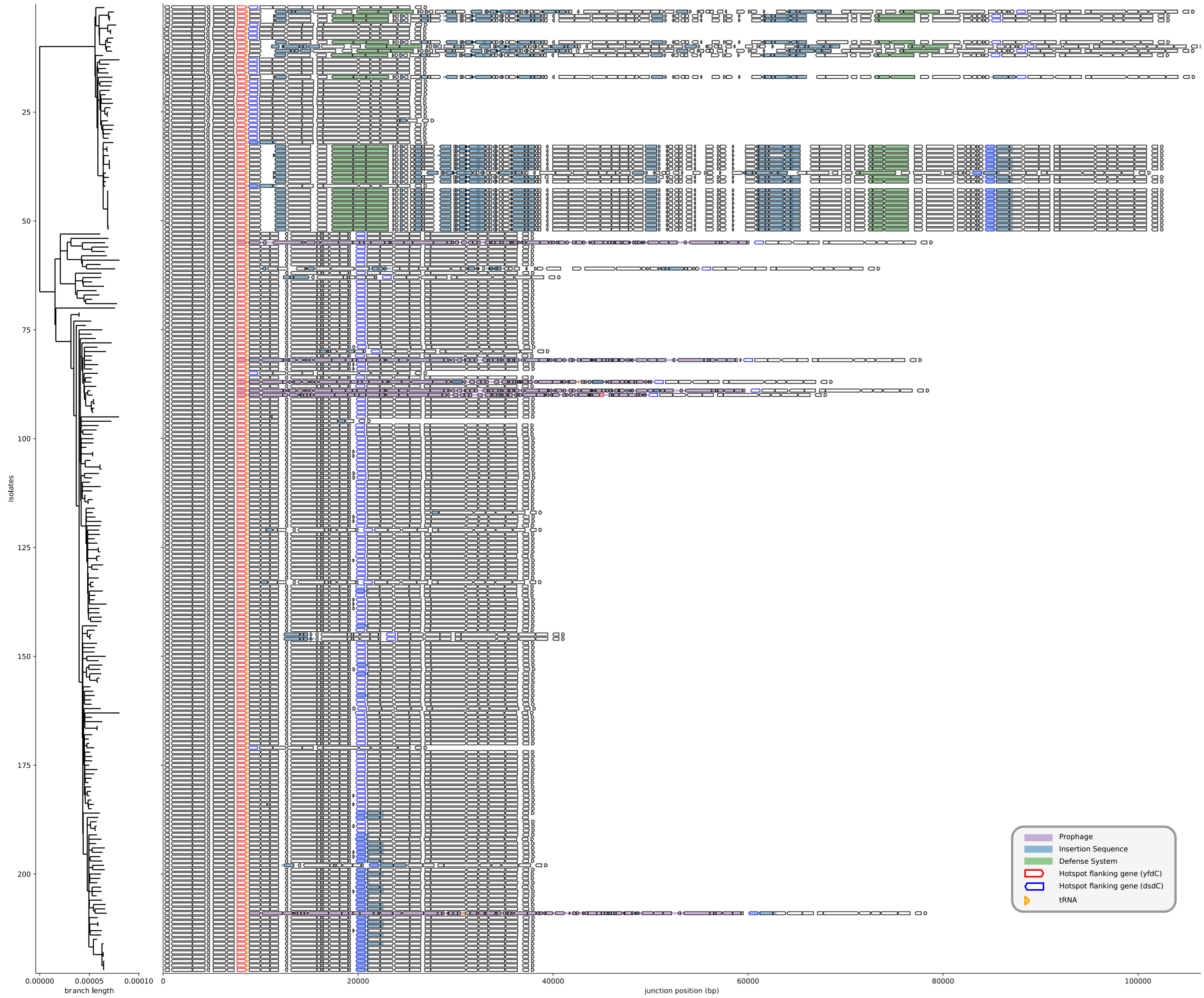


FIG. S20 **Example of functional content of complex junction containing hotspot 11**. Linear representation for the genomic region contained in a complex junction featuring 27 different distinct paths and a total pangenome length of 240 kbp. This junction contains the region named *hotspot 11* in (Hochhauser *et al.*, 2023), identifiable by its flanking genes (blue and red genes). Orange annotations are tRNA genes, while colored regions highlight annotations associated to prophages (purple), insertion sequences (blue) or defense systems (green).

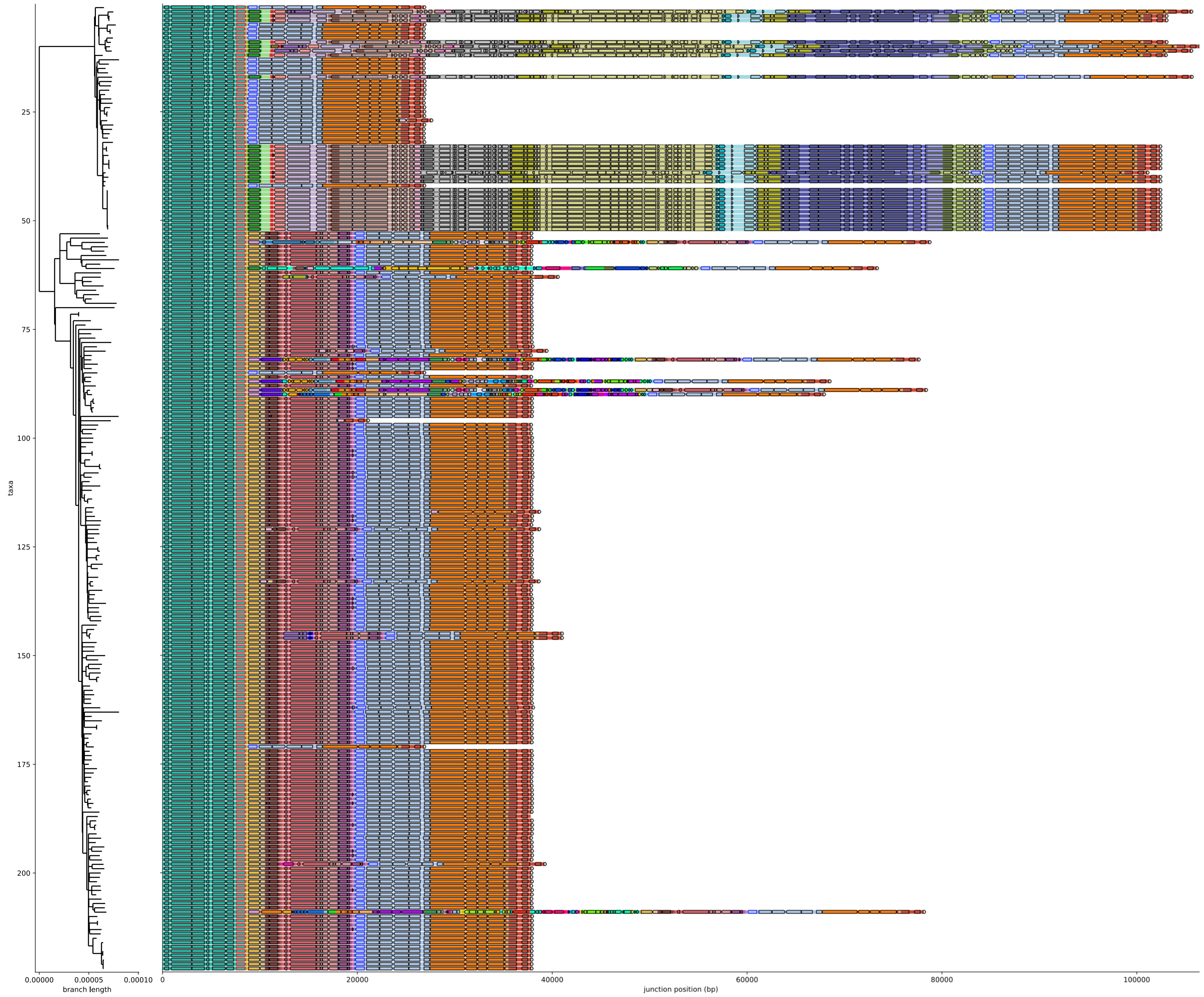


FIG. S21 **Example of structural diversity of a complex junction containing hotspot 11**. The figure contains the same junction depicted in fig. S20, with colors indicating regions belonging to different blocks in the pagenome graph representation. The first (teal) and last (red) block correspond to the flanking core blocks of the junction.

#### XI. COMPARISON TO GENE-BASED PANGENOME ANALYSIS USING PANX

To compare our results with gene-based methods we use PanX (Ding *et al.*, 2018) to group gene annotations in our dataset into orthologous gene clusters. PanX identifies 7863 gene clusters (GCs). The total detected pangenome size is of 6.85 Mbp, compatible with the one obtained by PanGraph (7.99 Mbp) considering that the latter also includes non-coding regions (on average 11.6% of the genome). PanX also produces a phylogenetic tree from the reconstructed core genome alignment, but it does not mitigate the effect of recombination. We find that indeed this tree is very similar to the one obtained with our method if we exclude the recombination filtering step. In general, the pangenome frequency distribution recovered by the two tools is highly compatible (see fig. S22). In more than 95% of cases, genes identified as core by PanX are completely included in a single core-block by PanGraph.

The main incompatibility between the two methods is found in *almost-core* genes: PanX provides an excess of genes found in all but one isolate. 40% of these almost-core genes (227/688) are placed by PanGraph in core-blocks, suggesting that homology is mostly maintained even in the isolate where the gene is inferred to be absent, and the absence of an annotation might be due to a frameshift or a premature stop codon. To confirm this hypothesis for each of the 227 almost-core genes assigned to core-blocks:

- We extract from the block alignment produced by PanGraph the region corresponding to the almost-core gene.
- For every entry in the alignment we detect whether it presents frame-shifts or premature stop codons.
- If a frame-shift or a premature stop codon is detected in exactly only the isolate where the gene annotation is missing, then we attribute the absence of the annotation to either of these causes.

In 67% of cases the absence of the annotation can be attributed to a frameshift, and in 28% of cases to a premature stop codon. Manual inspection of the remaining 5% of cases reveals frameshifts and stop codons also in these cases, but in multiple isolates. The fact that in some cases premature stop codons are close to the end of the sequence might conserve the annotation in these other isolates.

Overall, this shows that the pangenome graph obtained using PanGraph has a pangenome frequency compatible to the one obtained by traditional gene-based methods. Moreover, gene-based methods might lead to overestimation of the number of structural loss events for the reasons described above. By not relying on annotations, PanGraph can provide a more accurate estimate for the rate of large-scale gain/loss changes in the genome.

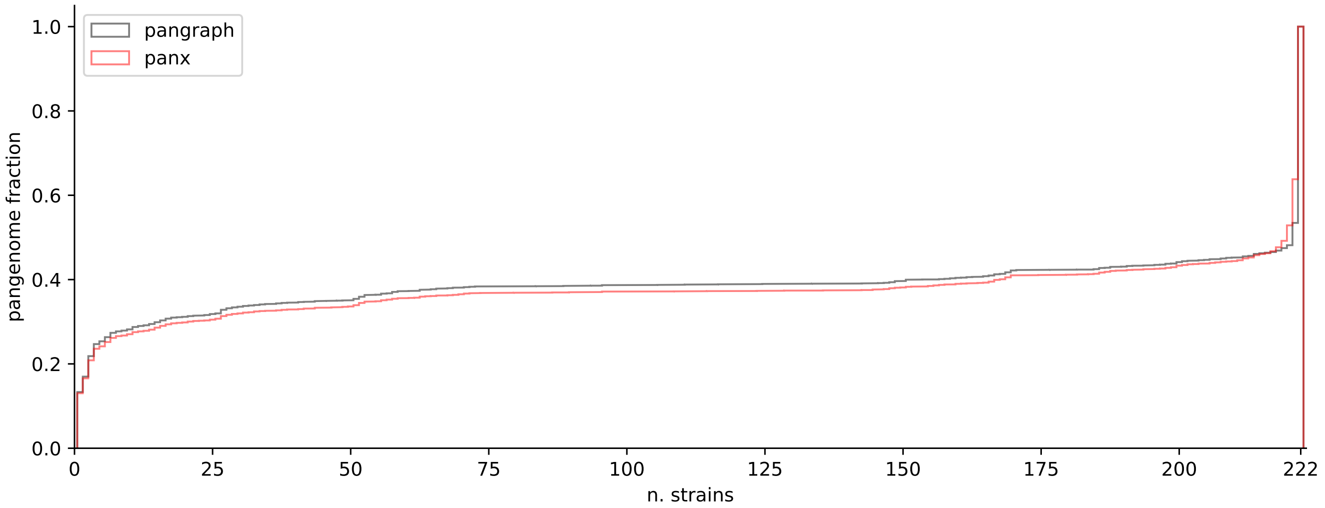


FIG. S22 **Comparison of pangenome frequency as estimated by PanX and PanGraph**, for the 222 isolates in our dataset. For each gene or block, we extract the number of isolates in which it is present, and plot the normalized cumulative distribution of these frequencies weighted by gene/block length.

#### XII. HETEROGENEITY IN RATES OF EVOLUTION OF GENOME STRUCTURE

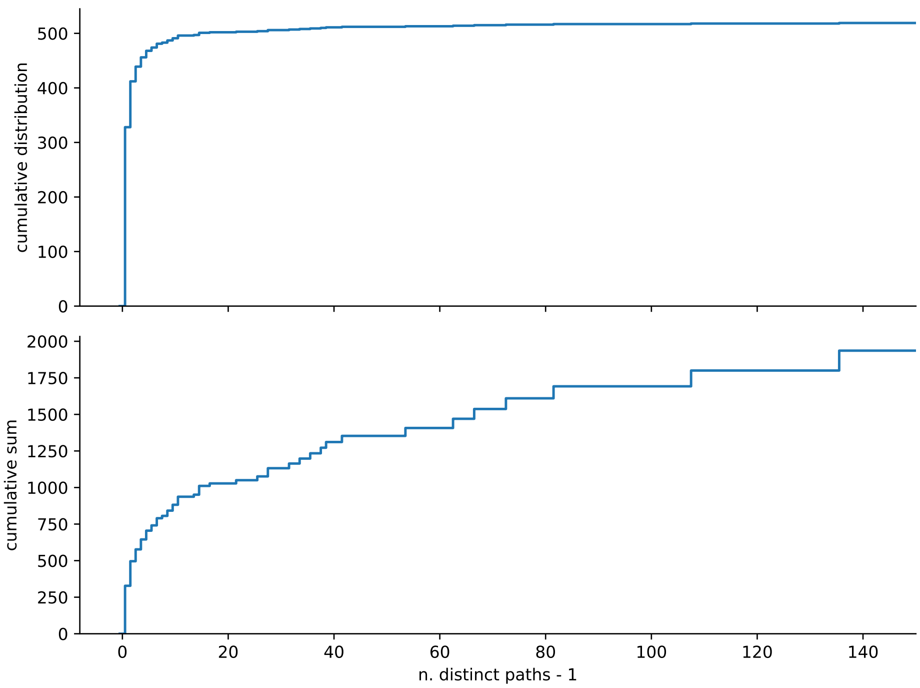


FIG. S23 **Quantification of minimal number of structural changes via amount of structural diversity in junctions.** We can obtain a lower bound for the total amount of structural changes that must have occurred in the evolutionary history of our dataset by considering that each junction with $n$ distinct paths must have been subject to at least $n-1$ structural changes. **Top**: cumulative distribution of $n-1$ values for all of the junctions in our dataset. **Bottom**: cumulative sum of $n-1$ values for all of the junctions in our dataset. This totals to 1936 events, corresponding to roughly one every 3 mutations on the (filtered) core genome alignment. Binary junctions contribute to roughly one-sixth of the total, and complex junctions with $n > 20$ to roughly half.
